# ApoE Lipidation State Directs Immunometabolic Reprogramming of Human Microglia

**DOI:** 10.64898/2026.05.04.722733

**Authors:** Tsion G. Shiferaw, Snigdha Sarkar, Katelyn M. Baker, Rowan S. Wooldridge, Hannah M. Binfet, Victoria N. Prozapas, Chinemerum P. Ogbu, Athena A. Schepmoes, Isaac K. Attah, Christy S. Niemeyer, Kayla G. Sprenger, James E. Hassell, Robert H. Eckel, John T. Melchior, Kimberley D. Bruce

**Affiliations:** Division of Endocrinology, Metabolism and Diabetes, University of Colorado Anschutz Medical Campus, Aurora, CO, USA; Biological Sciences Division, Earth and Biological Sciences Directorate, Pacific Northwest National Laboratory, Richland, WA, USA; Department of Neurology, University of Colorado Anschutz Medical Campus, Aurora, Colorado, USA; Department of Chemical and Biological Engineering, University of Colorado Boulder, 3415 Colorado Ave, Boulder, CO 80303; Department of Pathology and Laboratory Medicine, University of Cincinnati, Cincinnati, OH 45237; Department of Neurology, Oregon Health and Science University, Portland, Oregon 97239 USA

**Keywords:** Apolipoprotein E, APOE ε4, Microglia, Alzheimer’s disease, Lipid Metabolism, Holotomography

## Abstract

**Introduction:** ApoE4 is the strongest genetic risk factor for Alzheimer’s disease (AD). Emerging evidence suggests that ApoE4 increases AD risk by disrupting microglial metabolism and function. However, whether ApoE lipidation state contributes to microglial dysfunction remains poorly understood.

**Methods:** Human microglia were treated with lipid-free or lipid-bound ApoE3 or ApoE4. Label-free live-cell holotomography and global proteomics were used to assess isoform- and lipidation-specific effects on lipid droplet dynamics, mitochondrial morphology, and microglial phenotype.

**Results:** ApoE4 treatment resulted in fewer but enlarged lipid droplets and increased mitochondrial fragmentation compared to ApoE3, effects that were enhanced by lipid-bound ApoE4. Proteomic analyses revealed a strong type I interferon response in cells exposed to lipid-free ApoE, which was exacerbated by lipid-free ApoE4.

**Discussion:** These findings indicate that lipid-bound ApoE4 drives metabolic reprogramming, whereas lipid-free ApoE4 promotes inflammatory signaling, identifying ApoE lipidation as a critical modifier of ApoE4-associated AD risk.

## 1. Background

Alzheimer’s disease (AD) is a progressive age-associated and ultimately fatal neurodegenerative disease. Recent estimates predict that by 2050, 82 million people will be living with AD and AD-related dementias^1^. Pathological hallmarks of AD include accumulation of amyloid β (Aβ), tau tangles, neuroinflammation, lipid droplet accumulation, and mitochondrial dysfunction^2–4^. The ε4 variant of apolipoprotein E (ApoE4) is the strongest genetic risk factor for AD. Individuals homozygous for ApoE4 have up to a 15-fold increased risk of developing AD, compared to individuals carrying ApoE2 or ApoE3, which confer a protective or neutral risk for AD, respectively^1,5,6^. Alarmingly, current recent FDA approved therapeutics have limited benefit in ApoE4 carriers and may even worsen cognitive decline^7^, highlighting the need to better understand the cellular and molecular mechanisms by which ApoE4 drives AD neuropathogenesis.

Apolipoprotein E (ApoE) is the principal scaffold protein of brain lipoproteins, which mediate lipid transport throughout the central nervous system and facilitate horizontal lipid flux between neurons and glial cells^8,9^. These functions are enabled by ApoE’s distinct structural domains, including an N-terminal receptor-binding region and a C-terminal lipid-binding domain, which coordinate lipid and lipoprotein exchange^10^. ApoE exists in three main isoforms, which arise from two single-nucleotide polymorphisms (SNPs), rs429358 (C to T) and rs7412 (C to T), resulting in amino acid substitutions at positions 112 and 148^11,12^. Specifically, ApoE3 has a cysteine at 112 and arginine at 158; ApoE2 has a cysteine at 112 and 158; and ApoE4 has an arginine at both 112 and 158^11,12^. Notably, these minimal amino acid substitutions lead to structural differences have a major impact on receptor and lipid binding^12–14^. While lipidation-specific conformational changes remain an active area of investigation, ApoE4 is thought to be relatively lipid-poor, leading to immunometabolic reprogramming of microglia, which contributes to AD pathology^15^. However, the relatively low abundance of lipid-bound ApoE in the CNS, compared to circulating lipoproteins makes its isolation technically challenging, limiting our understanding of the lipidation- and isoform-specific mechanisms by which ApoE4 drives microglial dysfunction.

In the brain, ApoE is predominantly produced by astrocytes, although neurons, oligodendrocytes, and microglia also express and secrete ApoE^10,16^, particularly during development, damage, and disease^17^. Microglia, the brain’s resident innate immune effector cells, play a key role in protecting against the development of AD by detecting and clearing amyloid β (Aβ) plaques and tau tangles^16^. However, dysfunctional microglia are a major hallmark of AD and are characterized by impaired phagocytosis of protein aggregates, elevated neuroinflammatory signaling, lipid droplet (LD) accumulation, mitochondrial dysfunction, and glycolytic shifts, defined as immunometabolic reprogramming^18–21^. Given the potential to improve microglial function and AD outcomes, many recent studies have focused on identifying the factors that drive microglial dysfunction. Overwhelmingly, ApoE4 expression has been shown to promote microglial activation, mitochondrial impairment, inflammatory signaling, and LD accumulation^22–26^. However, due to the lack of available comparable sources of lipid-free and lipid-bound ApoE4, whether the lipidation status of ApoE4 drives microglial dysfunction—and the mechanisms underlying this dysfunction—remain unknown.

To address this gap, we generated and validated both lipid-free and lipid-bound ApoE^27^. As expected, lipid-free ApoE4 increased LD accumulation compared to ApoE3 in primary mouse microglia. Since typical imaging approaches rely on labeled lipids, which can alter baseline metabolism and preclude longitudinal analyses, here, we employed label-free live-cell holotomography to assess isoform- and lipidation-specific effects of ApoE on LD and mitochondrial dynamics. Using this approach, we found that supplementing human microglia with ApoE4 impairs cell health. Notably, lipid-bound ApoE4 reduced cellular ramification—a hallmark of homoeostatic microglia—and drove the accumulation of larger LDs. To probe mechanisms underlying LD expansion, we examined mitochondrial morphology. ApoE4 exposure led to fewer, shorter mitochondria, and lipid-bound ApoE4 further induced mitochondrial fragmentation, evidenced by shorter branch lengths. Proteomic analyses corroborated these findings, revealing downregulation of mitochondrial proteins and a pronounced type I interferon (IFN) response, which was strongest in cells treated with lipid-free ApoE4. Overall, these results demonstrate that ApoE4 exerts distinct immunometabolic effects depending on its lipidation state. Specifically, while lipid-bound ApoE4 drives metabolic dysfunction, lipid-free ApoE4 amplifies inflammatory signaling. These findings highlight the importance of considering lipidation status when evaluating ApoE4-mediated microglial dysfunction underlying AD neuropathogenesis.

## 2. Methods

### 2.1. Generation of lipid-free and lipid-bound ApoE isoforms

#### 2.1.1. Protein expression and purification

A pET32a expression vector, containing a thioredoxin fusion protein and mature human ApoE3 and ApoE4, was modified to introduce a cleavage site for tobacco etch virus (TEV) protease (ENLYFQ/X), as previously described for ApoA1^28^. The plasmids were transformed into *E. coli* BL21 cells, and transformants were cultured at 37°C in Luria-Bertani medium with ampicillin (100 µg/mL) to select transformants. Protein expression was induced using 0.1 mM isopropyl β-d-thiogalactopyranoside under shaking at 225 revolutions per minute (RPM) for 2 h at 37°C. Post-induction, cells were pelleted by centrifugation and stored at -40°C until ready for protein isolation. Cells were subsequently thawed on ice, resuspended in binding buffer (5 mM imidazole, 500 mM NaCl, 20 mM Tris-HCl, pH 7.9), and lysed at 4°C by probe sonication. Insoluble fractions were removed by centrifugation, and the soluble lysate was applied to His-bind columns (Novagen). Columns were washed extensively, and protein was eluted using elution buffer (1 M imidazole, 500 mM NaCl, 20 mM Tris-HCl, pH 7.9). The His-tag was cleaved with TEV protease at a protein-to-enzyme ratio of 20:1 for 2 h at room temperature. Cleaved samples were reapplied to His-binding resin to remove uncleaved protein. Purified protein fractions were pooled, dialyzed in 10 mM ammonium bicarbonate (pH 8.1), and lyophilized for storage at -80°C until ready for use. For experiments with lipid-free ApoE, the lyophilized protein was resolubilized in 3 M guanidine hydrochloride overnight and refolded in Standard Tris Buffer (STB; 10 mM Tris, 150 mM NaCl, 1 mM EDTA, 0.01% Na azide, pH 7.8). Protein was further purified in STB with no azide using size exclusion chromatography (SEC) with a Superdex 16/600 column. Fractions containing the pure protein were pooled, concentrated, and quantified using the modified Lowry assay.

#### 2.1.2. Generation of ApoE-containing reconstituted high-density lipoproteins (rHDL)

To generate ApoE-containing particles, lyophilized ApoE protein was unfolded, refolded, and purified by fast protein liquid chromatography in STB buffer, as described above. Reconstituted high-density lipoprotein (rHDL) particles were generated using the cholate dialysis method^29^ using molar ratios of 140:1 (1-palmitoyl-2-oleoyl-sn-glycero-3-phosphocholine (POPC: ApoE).):APOE). Briefly, micelles were generated through the addition of POPC (Avanti Polar Lipids, Cat: 850457C) and cholate (cholate:POPC ∼1.4). Prior to addition to micelles, ApoE protein was treated with 10 mM dithiothreitol (DTT) in STB and incubated at 37 °C for 1 h. Protein was incubated with micelles for 1 hour at 37°C to allow complexes to form. Cholate was removed by dialyzing particles into phosphate-buffered saline (1.9 mM KH_2_PO_4_, 8.1 mM Na_2_HPO_4_, 140 mM NaCl, 0.01% EDTA, 0.01% Na azide). All particles were further purified using size exclusion chromatography (SEC) and a single Superdex 200 10/300 column in STB with no azide. Where indicated, particles were labeled with 18:1 Liss Rhodamine PE (Liss-PE) (Avanti, Cat: 384833-00-5), as previously described3230. Briefly, Liss-PE was solubilized in dimethyl sulfoxide (0.01 mg/mL). 1 mg of particles were incubated with 15 μL of label overnight at room temperature. Particles were purified by SEC, and traces were monitored by both ultraviolet (560 nm) and fluorescence (583 nm). Fluorescence was monitored in real-time as particles eluted from the column using a Dionex Ultimate 300 (Thermo Fisher Scientific) equipped with a flow cell integrated between the columns and fraction collector. Signal was acquired using an excitation wavelength of 560 nm and emission wavelength of 583 nm. Fractions corresponding to the pure particles were collected, aliquoted, snap-frozen in liquid nitrogen and stored at -80°C until ready for use.

### 2.2. Cell culture

#### 2.2.1. Human fetal microglia (HMg) cell maintenance

Human fetal microglia (HMg, HMC3) cells were cultured in MEM/EBBS (Hyclone, Cat: SH30024.02 Lot: AK30808187) and supplemented with 10% fetal bovine serum (FBS; Benchmark™ Cat: 100-106) plus 1% penicillin/streptomycin (Corning, Ref: 30-002-CI) and 1% Sodium Pyruvate (only in MEM/EBBS; Gibco, Ref: 11360-070, Lot: 2192805). The cells were maintained at 37 °C with 5% CO_2_ and 95% humidity. Cells were passaged at >90% confluency. These media formulations were used exclusively for cell maintenance; treatment conditions are described separately. As a critical control, genomic DNA was extracted from HMg cells and ApoE genotype was determined using allele specific qPCR-based genotyping (CD genomics)^30^. HMg cells are homozygous for *ApoE3* and are male.

#### 2.2.2. Primary microglia culture and ApoE treatment

Primary murine microglia were isolated from mixed glial cultures, as previously described^31^. Isolated microglia were cultured in Dulbecco’s Modified Eagle’s medium (DMEM)/Ham’s F-12 medium (Corning), supplemented with 10% FBS (Benchmark™) and 1% penicillin-streptomycin (Corning). Upon reaching confluency, cells were transferred (seeding density of 130,00cells/well) to Poly-D-lysine (Gibco, Ref: A38904-01) coated 24-well plates (IbIDI, Lot: 240909) and treated with fluorescently labeled ApoE-containing rHDL (concentration of rHDL-E3 at 0.05 mg/mL rHDL-E4 at 0.04 mg/mL) or lipid-free ApoE (concentration of LF-E3 at 0.03 mg/mL and LF-E4 at 0.02 mg/mL), in order to maintain a constant lipid concentration of 200 μM. ApoE treatment experiments were performed in DMEM/Ham’s F-12 medium (Corning), supplemented with 1% FBS (Benchmark™) and 1% penicillin-streptomycin (Corning) for 6 hours.

### 2.3. Fluorescence imaging of primary microglia

Following ApoE treatment, primary microglia were washed with 1% fatty acid-free bovine serum albumin (BSA) in phosphate-buffered saline (PBS) (Cytiva, Lot: AL30843974) to remove unbound proteins and lipids. Cells were then fixed with 4% paraformaldehyde (PFA) for 10 min at room temperature. After fixation, PFA was removed, and cells were stored in PBS overnight at 4°C. Cells supplemented with ApoE-containing rHDL were stained with Alexa Fluor 488-conjugated phalloidin (1:1,000) (Invitrogen, Cat: R415) to visualize F-actin and DAPI (1:500) (Invitrogen, Lot: 2942291) to label nuclei. For cells treated with lipid-free ApoE, Alexa Fluor 620-conjugated phalloidin (1:1,000) (Invitrogen, Cat: AB176757) was used to visualize F-actin, BODIPY (1:1,000) (Invitrogen, Lot: 2201649) to label neutral lipids, and 4′,6-diamidino-2-phenylindole (DAPI) (1:500). After staining, cells were washed with PBS and imaged using an Olympus XM10 inverted microscope (Evident Corporation, Nagano, Japan) at 40X magnification (0.60NA, model LUCPLFLN40X, Evident Corporation, Nagano, Japan). Images were acquired using the Olympus cellSens Software (version 1.18, Evident Corporation, Nagano, Japan). Due to differences in staining procedures between the treatment groups, exposure times varied.

#### 2.3.1. Primary Microglia Image Analysis

Images were processed using FIJI^32^. Cell clusters were manually segmented. Following segmentation, the ‘Analyze Particles’ tool was used to identify regions of interest with a minimum pixel length of 200 to select for cells and exclude debris. Regions of interest were applied to a DAPI image that was converted to binary with the ‘Li dark’ threshold setting, and the maximum pixel value was measured. Regions with a max DAPI value of 0 (phalloidin positive, DAPI negative) were later excluded from final analysis. On the images from cells treated with fluorescently labeled rHDL, background signal and noise were mediated by setting a minimum pixel value of 205, a maximum value of 555, and then using the rolling ball background subtraction tool (radius 25 pixels). Rolling-ball background subtraction was also applied to the lipid-free ApoE images. Using the corresponding regions of interest, mean fluorescent intensity of BODIPY or 18:1 Liss-PE per cell was collected. Prior to statistical analysis, DAPI-negative objects were excluded using R^33^ with dplyr^34^ and tidyr^35^.

### 2.4. Label-Free, Live-Cell holotomography

#### 2.4.1. Cell culturing and ApoE treatment

HMg cells (passage 4, P4) were cultured to confluency in T75 flasks (Falcon, Lot: 353135). Cells were subsequently cold seeded into a 96-well microplate with a glass bottom (Nanolive, Lot: 053585) at a density of 10,000 cells per well and incubated overnight at 37 °C in a humidified 5% CO_2_ to allow for attachment. The following day, the medium was aspirated and replaced with treatment media consisting of either ApoE-containing rHDL or lipid-free ApoE at a final concentration of 0.01 mg/mL. Treatments were prepared in DMEM High glucose (Hyclone) supplemented with 1% lipoprotein-depleted FBS (Lipid Free FBS: Kalen Biomedical, B5148301). Each condition was imaged at two time points (4-6 hours and 22-24 hours).

#### 2.4.2. Holotomography image acquisition and analysis

Live cells were positioned in a stage-CO_2_ incubator within the holotomography microscope (Nanolive, 3D Cell Explorer-96focus light microscope). The incubator was maintained at 37 °C, 5% CO_2_, and 100% humidity, providing optimal cell maintenance conditions throughout image acquisition. Images were acquired 4-6 or 20-22 h post-ApoE treatment, using the same wells each time (n = 4 wells per condition). The machine is equipped with a rotating arm that rotates the object beam 360° around the sample and was set to take 5×5 grid scans of each condition, approximately every 20 minutes, for 10 cycles within the given time frame. Image slices were taken in the z-stack, covering 30 µm in depth, and reconstructed into maximum-intensity projections (MIPs). All post-analysis was completed on the Eve Explorer software (Nanolive), which calculates various metrics of cell morphology, LDs (Smart Lipid Droplet Assay ^LIVE^), and mitochondria dynamics (Mitochondrial Assay) for each cell and each experimental group. A detailed description of each metric interpretation is shown in **Supplementary Table 1**. In all the analysis outcomes for LDs and mitochondria shown in this manuscript, the data are presented as per-cell averages within each condition.

### 2.5. Proteomics

#### 2.5.1. Culturing and treatment

As previously described, P4 HMg cells were grown in T75 flasks to confluency. Cells were then plated in a 6 well plate at 250,000 cells/well in MEM/EBBS (Hyclone, Cat: SH30024.02 Lot: AK30808187) overnight to attach. The following morning, the media was removed and treated with ApoE-containing rHDL or lipid-free ApoE at 0.01 mg/mL in DMEM High glucose (Hyclone, Ak30769531) supplemented with 1% lipoprotein-depleted FBS (Lipid Free FBS: Kalen Biomedical, B5148301), for 6 or 24 hours.

#### 2.5.2. Cell harvest for proteomics

To harvest the cells, the media was removed, and the plate was washed twice with warmed PBS (Cytiva, AL30843974). To avoid cell death from mechanical scraping, cells were trypsinized (0.25x) for 5 min. Trypsin reaction was stopped with complete media (DMEM High glucose). Cells were transferred to a 15 mL Falcon tube and spun at 500 g for 5 minutes. The media was aspirated, and the cell pellet was resuspended in PBS and transferred to an Eppendorf tube to spin down again (500 g for 5 min), aspirated, and resuspended in PBS. After the final aspiration, the cell pellet was snap-frozen in liquid nitrogen and stored at -80 ℃ until further analysis.

#### 2.5.3. Sample preparation for proteomics analysis

Cells were thawed on ice and lysed by resuspending the pellets in 8 M urea solution. The denatured proteins in the cell lysates were reduced by incubating with DTT, (5 mM) at 60 °C for 30 min with shaking at 500 rpm. Post-reduction, samples were alkylated by treating with iodoacetamide (40 mM) at 60 °C for 30 min. The protein extracts were digested, first with LysC at an enzyme:substrate ratio of 1:100 for 2 h at 25 °C, and then with trypsin at an enzyme: substrate ratio of 1:50 for 3 h at 37 °C with constant shaking at 500 rpm. The digestion reaction was quenched with 0.1% formic acid, and the resulting peptides were desalted using C18 solid phase extraction. The final concentrations were determined using a Pierce BCA protein assay (ThermoFisher Scientific, Cat: A55864). The peptide mixtures were diluted to 0.1 mg/mL in 0.1% formic acid and stored at -20 °C until ready for analysis by liquid chromatography-mass spectrometry (LC-MS/MS).

#### 2.5.4. Proteomics data acquisition

The peptide mixtures were analyzed using data-independent acquisition (DIA)^36^. LC-MS/MS analysis was performed using a Thermo Scientific Q Exactive HF-X mass spectrometer coupled to a Dionex Ultimate 3000 chromatography setup, as previously described^37^. Briefly, the peptide mixtures were separated on a reverse-phase C18 column (30 cm × 75 μm, 1.7 μm Waters Acquity BEH particles) over a 2 h gradient at 200 nL/min using two buffers: (1) 0.1% formic acid in water, and (2) 0.1% formic acid in acetonitrile. For DIA, MS1 scans were acquired at a resolution of 60k over a mass range of 400–900 m/z, with an AGC target of 1×10⁶ ions and a maximum injection time of 128 ms. HCD fragmentation was performed with a normalized collision energy of 30. MS2 spectra were collected at a resolution of 30k across the same 400–900 m/z range, using 10 Da isolation windows, an AGC target of 1×10⁶, and a maximum injection time of 64 ms. MSX isochronous injection times were enabled, with the MSX count set to 1.

#### 2.5.5. Proteomics data analysis

The .raw files generated by the Thermo mass spectrometer were converted to. mzML using MSConvert (ver 3.0.24239.0) using the peakPicking option. The .mzML files were searched using FragPipe (ver 22.0) equipped with MSFragger (ver 4.1) and DIANN (ver 1.8.2), with the ‘DIA_SpecLib_Quant’ workflow. The searches were performed against the *Homo Sapiens* UniProt/SwissProt database (Proteome ID #UP000005640, 20,417 entries, downloaded 11/14/2025), supplemented with common contaminants and decoy. The following parameters were used: (1) enzyme specificity was selected as strict-trypsin, (2) allowed missed cleavages was set to 4, (3) precursor mass tolerance was set to ±10 ppm, (4) fragment mass tolerance was set to 20 ppm, (5) variable modifications─N-terminal peptide acetylation, oxidation at methionine, (6) fixed modification─carbamidomethylation at cysteine, and (7) the False Discovery Rate (FDR) was set to 0.01. The protein-level output from DIANN was further analyzed using the R package RomicsProcessor (v1.1, https://doi.org/10.5281/zenodo.10459578). Briefly, protein intensities were log2-transformed, filtered to allow maximum missingness of 40% within each group, and the remaining missing values (<0.5%) were imputed following a previously described method^38^.

#### 2.5.6. Identification of Proteins of Interest, Differentially Abundant Proteins (DAPs), and pathway analysis

An unpaired two-sample *t*-test and Wilcoxon rank-sum test were performed on all proteins to determine differences between groups. Proteins with a p-value of less than 0.05 were referred to as proteins of interest (POI). Proteins with a p-value of less than 0.05 and an adjusted p-value of less than 0.1 reached the threshold of significance and were considered and referred to as differentially abundant proteins (DAPs). MetaboAnalyst 6.0 was used to visualize the proteins of interest using PLS-DA, VIP scores, heatmaps, and volcano plots, as previously described^39,40^. Pathway analysis was performed using STRING 12.0^41^ using the significantly upregulated and downregulated DAPs to identify upregulated and downregulated ‘KEGG’ and ‘biological processes’ pathways. Pathways with identical protein profiles were combined and titles were shortened to reduce space.

### 2.6. Quantitative-PCR

Total RNA was isolated from HMg cells using the Trizol-chloroform extraction method^42^. qScript cDNA synthesis kit (Quantabio, Cat: 95047) was used to synthesize cDNA from RNA. Gene expression levels relative to the reference gene (UBC) were quantified by qPCR, using the 2 (-ΔΔC(T)) method with lowest value normalization^43^. NCBI’s primer-blast tool (https://www.ncbi.nlm.nih.gov/tools/primer-blast) was used to design primers to human *SP100* and *STAT1* that crossed exon boundaries when possible (*SP100* Fwd: CATGCAGGAATACCCCGATTT; *SP100* Rev: GAGTTTTCACCAGTTCCTTGTTC. *STAT1* Fwd: CAAGTTCGGCAGCAGCTTAA; *STAT1* Rev: CACCACAAACGAGCTCTGAAT. *UBC* Fwd: FACTCTGCACTTGGTCGTCC; *UBC* Rev: GAATGCAACAACTTTATTGAAAGAA). Using a StepOnePlus instrument and software v2.3 (Applied Biosystems, Foster City, CA, USA), thermal cycling conditions for all qPCR studies were as follows: initial temperature of 50 °C for 2 min, 95 °C for 10 min, then 40 cycles of 95 °C for 15 s and 60 °C for 1 min.

### 2.7. Statistical analysis

When two groups were analyzed (e.g., primary murine fluorescence imaging), a nonparametric t-test (Mann-Whitney) was used to assess differences between groups. Wherein more than two groups were incorporated into the experiment (live cell imaging and proteomics) and analyzed a non-parametric (Kruskal-Wallis) was employed followed by a Tukey’s multiple test for multiple comparisons or a one-way analysis of variance (ANOVA) followed by the use of Dunn’s test for multiple comparisons, unless otherwise stated. Analysis was performed using GraphPad Prism Version 10.4.2 for Windows (GraphPad Software, Boston, Massachusetts, USA; www.graphpad.com). Results are presented as either violin plots with medians and quartiles, box plots with mean ± standard error of the mean (SEM), volcano plots with x-axis representing log_2_FC and y-axis representing -log_10_(p-value), heatmaps of proteins of interest. p<0.05 indicate significance, unless otherwise stated.

## 3. Results

### 3.1. Generation and characterization of lipidated and lipid-free ApoE3 and ApoE4

Although many studies have linked ApoE4 expression to microglial dysfunction, how human microglia differentially respond to ApoE3 and ApoE4 in their lipid-free and lipid-bound states has not been determined^44,45^. To address this, lipid-free ApoE3 and ApoE4 (LF-E3 and LF-E4, respectively) and reconstituted high-density lipoprotein-like lipoparticles (rHDLs) containing ApoE3 or ApoE4 (rHDL-E3 and rHDL-E4, respectively) were generated. To ensure accurate comparison between isoforms, rigorous preparation and quality control measures were performed on both the ApoE proteins and ApoE-containing rHDLs. ApoE protein was extensively purified by unfolding in guanidine-HCl, refolding, and polishing through SEC to ensure removal of any residual bacterial contaminants. Protein purity was >98% as assessed by SDS-PAGE analysis, showing single bands in each fraction collected from SEC for each isoform (**Supplementary Fig. 1**).

To test the impact of ApoE lipidation state, rHDL particles were generated containing either ApoE3 or ApoE4 from the same purified lipid-free ApoE protein stocks. Particles were generated using the same lipid-to-protein ratio to ensure that the primary variable between particles was limited to the ApoE isoform only. Particles were highly homogeneous, each contained two molecules of ApoE, and exhibited no differences in size as determined by SEC (**Figure 1A**). Homogeneity and particle sizes was further evaluated on pooled fractions using native gel electrophoresis, which confirmed that there were no major structural differences between particles containing ApoE3 or ApoE4 (**Figure 1B**). Similarly, quantification of phospholipids using a standard assay kit (Wako) revealed no significant differences in lipid content (data not shown). To assess the effects of fluorescent labeling overnight, particles containing Liss-PE were compared to non-labeled particles without incubation. SEC analysis revealed no observable differences in the UV absorbance traces from non-incubated samples, indicating that particle stability and homogeneity were maintained after labeling, and were suitable for the downstream analysis and cellular assays.

**Figure 1.**
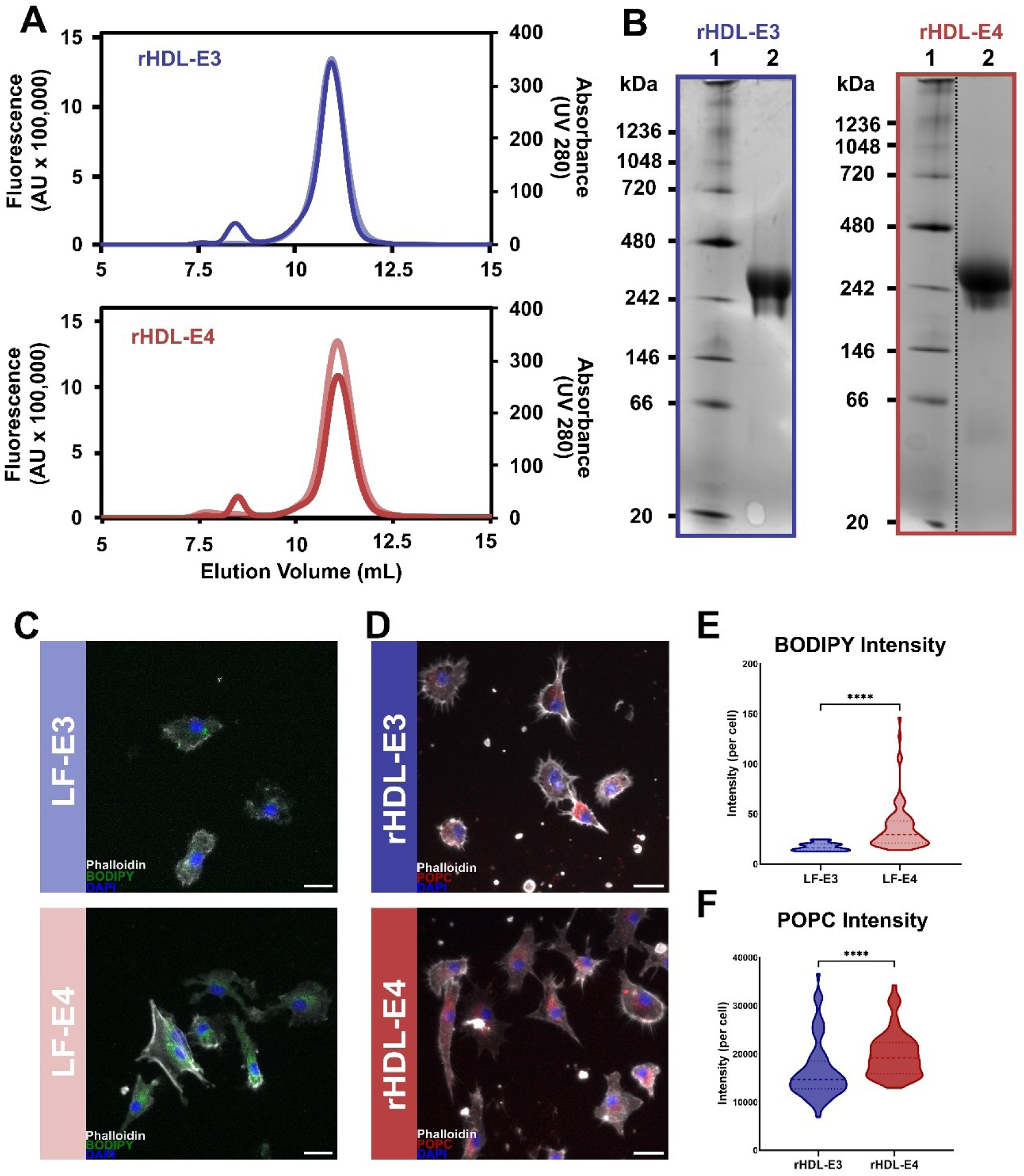
Lipid-free ApoE4 leads to increased lipid-droplet accumulation, whereas reconstituted HDL-like particles (rHDL) lead to enhanced lipid (POPC) uptake by primary microglia. **A.** Purity and homogeneity of ApoE-containing reconstituted high-density lipoprotein (rHDL) particles were evaluated by size exclusion chromatography (SEC) using a fluorescence lipoprotein detector (FLD) and a Superdex 200 10/300 column. For particles labeled with Liss-PE, fluorescence was monitored with an emission wavelength of 583 nm (dark line). Non-labeled particles were monitored by absorbance at 280 nm (lighter line). **B.** Fractions eluting between 10–12 mL were collected, pooled, and analyzed by native gel electrophoresis on an 4-16% gradient gel. Each lane contains 6 µg of protein. Protein on particles were visualized by Coomassie Blue staining**. C.** Fluorescence images of microglia treated with lipid-free ApoE3 (LF-E3) or lipid-free ApoE4 (LF-E4) (bottom). Cells were stained with Phalloidin (white, F-actin cytoskeleton), BODIPY (green, neutral lipids), and DAPI (blue, nuclei). Scale bar represents 100 µm. **D.** Fluorescence images of microglia treated with rHDLs containing fluorescently labeled POPC and ApoE3 (top) or ApoE4 (bottom). POPC (red) was quantified and cells were stained with Phalloidin (white), and DAPI (blue). Scale bar represents 100 µm. **E.** Quantification of lipid accumulation (BODIPY) shown as mean fluorescence intensity per cell. L-E4 treated cells showed higher BODIPY intensity compared to LF-E3 **F.** Quantification of POPC uptake shown as mean fluorescent intensity per cell. Cells treated with rHDL-E4 showed higher POPC internalization compared to cells treated with rHDL-E3. Data expressed as violin plots showing median, upper, and lower quartiles n=(3) *(****p < .0001*)

### 3.2. Lipid-free ApoE4 leads to increased lipid accumulation, and lipid-bound ApoE4 leads to enhanced lipid uptake by primary murine microglia

To determine how ApoE lipidation state and isoform influence microglial responses to ApoE, we treated primary murine microglia with either LF-E3 or LF-E4, or fluorescently labeled rHDL-E3 or rHDL-E4. For cells treated with LF-E3 or LF-E4, we measured neutral lipid accumulation after supplementation by quantifying intracellular BODIPY staining (Figure 1C, E). The rHDL-E3 and rHDL-E4 particles contained rhodamine-labeled 18:1 Liss-PE, allowing us to track particle uptake—or at least the uptake of particle-associated phospholipids—by quantifying fluorescent intensity per cell (Figure 1D, F). Cells treated with LF-E4 showed greater BODIPY intensity than those treated with LF-E3 (Figure 1C, E), consistent with reports that ApoE4 treatment or expression leads to increased microglial lipid accumulation^46^. We also observed that primary microglia treated with rHDL-E4 showed increased Liss-PE intensity (Figure 1D, F) compared to cells treated with rHDL-E3, suggesting that lipid-bound ApoE4, and its associated lipids, are taken up more readily than lipid-bound ApoE3. Although unexpected, it is consistent with recent studies supporting stronger interactions between ApoE4 and microglial lipoprotein receptors^47,48^.

### 3.3. Label-free, live-cell holotomography reveals reduced cell viability and cell size in human microglia treated with ApoE4

Recognizing the limitations of static measurements of lipids in live cells, we next employed holotomography microscopy, which enables live-cell, label-free imaging of cell morphology, LDs, and mitochondrial dynamics in real time^49–51^. In addition, given the volume of cells required for these assays and subsequent biochemical and proteomic assessments, we pivoted to using human fetal microglia (HMg) cells. Importantly, we used PCR-based sequencing to determine for the first time, that these male cells are homozygous for *ApoE3* (*ε3/ε3*). We performed morphological characterization of HMg cells treated with LF-E3, LF-E4, rHDL-E3, or rHDL-E4 at 6 h (**Figure 2A-F**) and 24 h (**Figure 2G-L**) post-treatment. Initial visualizations of the holotomography images are shown as maximum intensity projections (MIPs) (**Figure 2A and 2G**), which were then used to perform cell masking and segmentation (**Figure 2B and 2H**). Since metrics are dependent on the appropriate segmentation of cells and identification of cell boundaries from the subsequent cell masks, the accuracy of cell segmentation was carefully evaluated for each condition and time point. At 6 h, cells treated with LF-E3 or rHDL-E3 showed a more favorable health index (**Figure 2C**) compared to cells treated with LF-E4 or rHDL-E4. While cells treated with LF-E3 or rHDL-E3 exhibited similar morphologies, rHDL-E3-treated cells exhibited a modest increase in cell area compared to ApoE4 treated cells (**Figure 2D**). Cells treated with rHDL-E3 (**Figure 2F**) also showed increased ramification, compared to LF-E3, which is typically associated with microglial homeostasis.

**Figure 2.**
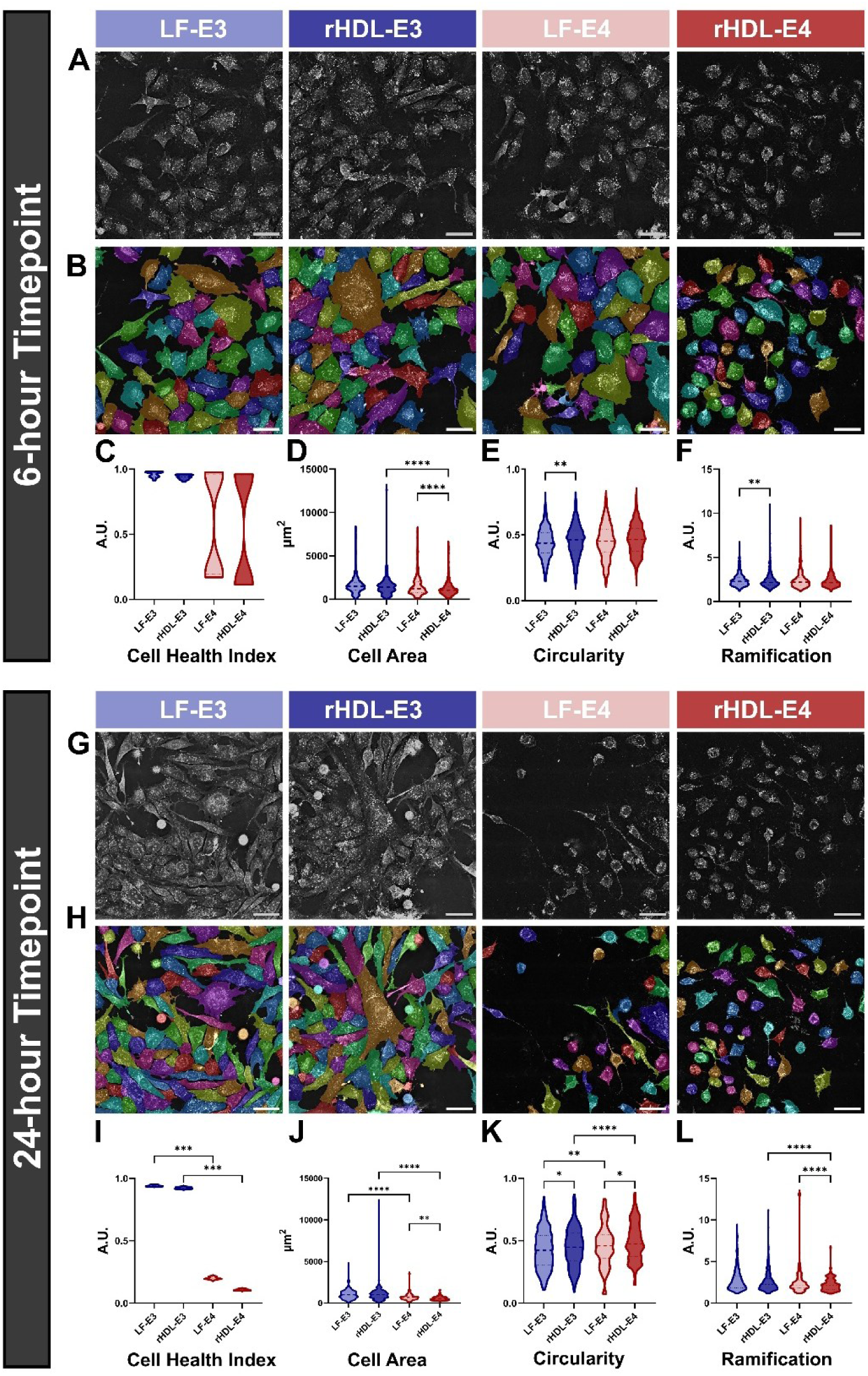
ApoE4 treatment increasingly reduces microglial viability and size over 24 hrs. Morphological characterization of human fetal microglia (HMg) treated with lipid-free (LF-E3, LF-E4) or lipid-bound ApoE3 and ApoE4 (rHDL-E3, rHDL-E4) at 6 h (**A-F**) and 24 h (**G-L**). **A.** Maximum intensity projections (MIPs) of the cells at 6 h. **B.** Cell segmentation at 6 h. **C.** Cell health index metric (6 h). **D.** Cell area metric (6 h). **E.** Circularity metric (6 h). **F.** Ramification metric (6 h). **G.** MIP of the cells at 24 h. **H.** Cell segmentation at 24 h. **I.** Cell health index metric (24 h). **J.** Cell area metric (24 h). **K.** Circularity metric (24 h). **L.** Ramification metric (24 h). Violin plots show median, upper, and lower quartiles (n=3 image, 50-200 cells per condition), *\* p<0.05, **p<0.01, ***p<0.001, ****p<.0001.* All scale bars represent 50 µm.

By 24 h, cells treated with LF-E3 or rHDL-E3 maintained a higher cell health index than cells exposed to LF-E4 or rHDL-E4 (**Figure 2I, Supplementary Table 1**). Cells treated with LF-E3 or rHDL-E3 were also larger than cells treated with LF-E4 or rHDL-E4 at this time point (**Figure 2J**). Furthermore, by 24 hrs, treatment with LF-E4 or rHDL-E4 resulted in increased circularity, suggesting transition to a more amoeboid and dysfunctional state in response to ApoE4. Consistent with increased microglial dysfunction, cells treated with rHDL-E4 became less ramified (**Figure 2L**). Taken together, this data suggests that while ApoE4 impairs microglial viability and morphology, the lipidation state of ApoE further modulates these microglial responses.

### 3.4. ApoE4 promotes the formation of larger, more densely packed lipid droplets

Microglial expression of ApoE4 has been repeatedly linked to increased and potentially pathogenic LD accumulation^22,52^. Despite this, the impact of exogenous lipid-bound and lipid-free ApoE on microglial LD dynamics has not been determined. To address this, we utilized holotomography to measure LD dynamics in live cells without the need for dyes or labels. HMg cells were treated with LF-E3, LF-E4, rHDL-E3, or rHDL-E4 for 24 h, with image and LD analyses performed at 6 and 24 h. LD segmentation and masking were performed on each individual LD within each cell, revealing abundant LDs in all conditions (**Figure 3A-B**).

**Figure 3.**
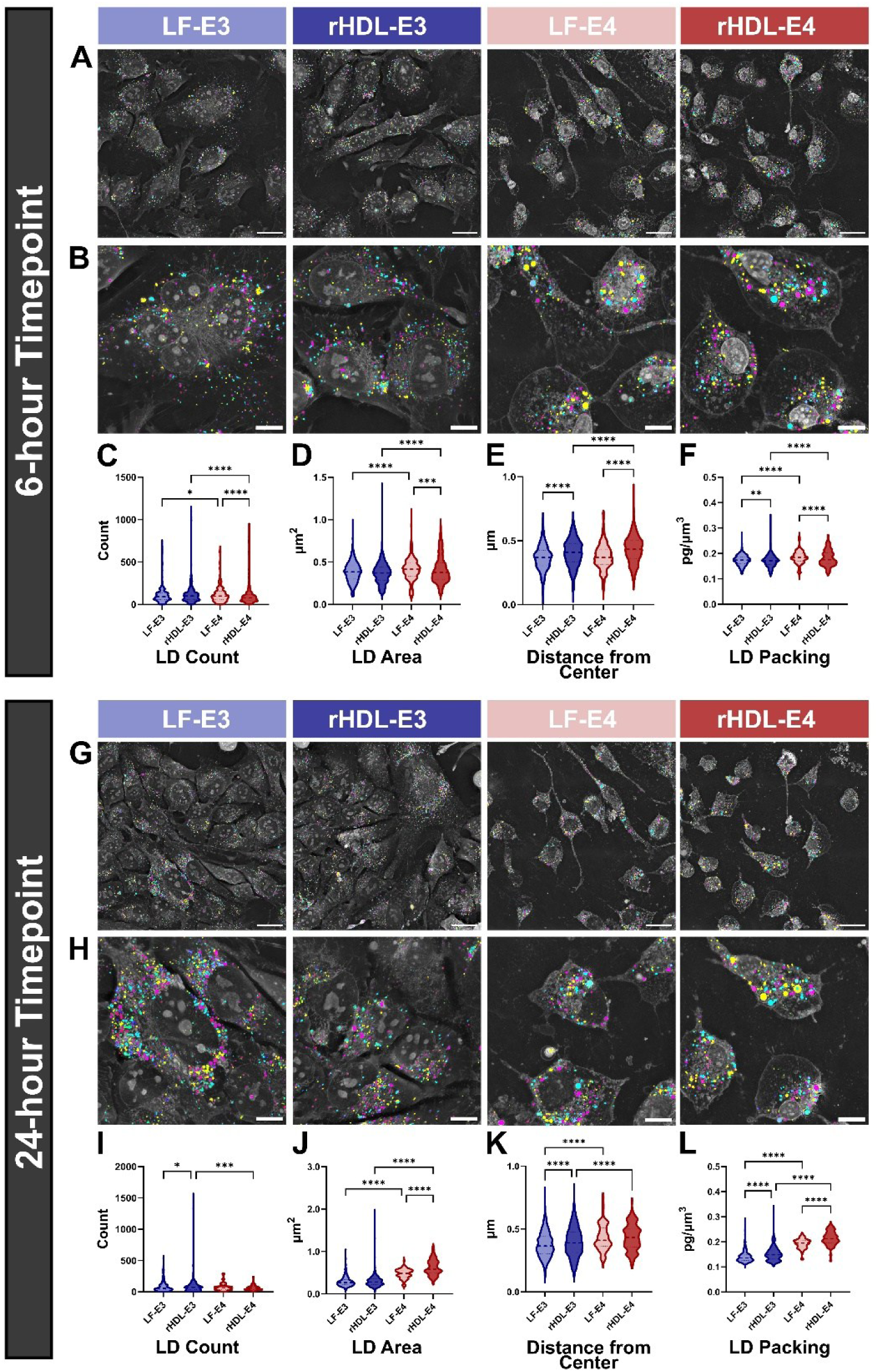
ApoE4 promotes the formation of larger, more densely packed lipid droplets. Lipid droplet (LD) characterization of human microglia treated with lipid-free (LF-E3, LF-E4) or lipid-bound ApoE3 or ApoE4 (rHDL-E3, rHDL-E4) for 6 h (**A-F**) and 24 h (**G-L**). **A.** Representative LD masks at 6 h. Scale bars represent 25 µm. **B.** Example, Zoomed-in LD mask at 6 h from images in Panel A. Scale bars represent 10 µm. **C.** LD count per cell (6 h). **D.** LD area per cell (6 h). **E.** LD distance from center (6 h). **F.** LD packing per cell (6 h). **G.** Representative LD masks at 24 h zoomed out. Scale bars represent 25 µm. **H.** Example LD mask at 24 h from images in Panel G. Scale bars represent 10 µm. **I.** LD count per cell (24 h). **J.** LD area per cell (24 h). **K.** LD distance from center (24 h). **L.** LD packing per cell (24 h). Violin plots show median, upper, and lower quartiles (n=3, 50-100 cells). *\*p<0.01, **p<0.001, ***p<0.0001, ****p<0.00001*.

By 6 h, LD counts were higher in rHDL-E3- and rHDL-E4-treated cells, suggesting that increased lipid supply may initially promote LD accumulation (**Figure 3C**). Given that these experiments were performed with de-lipidated media—to limit exogenous lipid and ApoE delivery to our defined source of ApoE—lipid supply is indeed greater in the rHDL-E3 and rHDL-E4 conditions. Notably, by 6 h, LD area was greatest in the LF-E4-treated cells (**Figure 3D**). In addition, LDs were further from the center in rHDL-E3- and rHDL-E4-treated cells (**Figure 3E**), indicating that lipid supply modifies LD processing and dynamics. Finally, we also noted that LDs were more densely packed when cells were exposed to LF-E4 (**Figure 3F**), suggesting increased protein content and potentially reduced lipolysis^53^.

24 h after ApoE treatment, LDs in the LF-E3- and rHDL-E3 treated cells were visibly smaller than LDs in the LF-E4- and rHDL-E4-treated cells (**Figure 3G-H**). This observation is consistent with quantification of LD area, which shows that cells treated with LF-E4 or rHDL-E4 exhibited the largest LDs (**Figure 3J**), implying ApoE4-specific changes in lipid mobilization, lipolysis, expansion, and/or biogenesis. LD size was further amplified when the cells were treated with rHDL-E4 versus LF-E4 (**Figure 3J**), suggesting a synergistic impact of ApoE4 and increased lipid supply on microglial LD expansion. In addition, LDs were further from the center and more densely packed when the cells were treated with LF-E4 or rHDL-E4, with the impact of ApoE4 being exacerbated in the presence of lipid (**Figure 3K-L**). Overall, this data suggests that ApoE4 leads to larger, more densely packed LDs by 24 h, an effect which appears to be amplified when ApoE4 is associated with the rHDL particle, increasing lipid supply.

### 3.5. ApoE4 promotes mitochondrial fragmentation

The accumulation of larger LDs in HMg cells treated with exogenous ApoE4 could be a consequence of other ApoE4-mediated bioenergetic changes within the cell. To address this, we employed mitochondrial analysis following label-free, live-cell holotomography imaging at both 6 and 24 h. Individual mitochondria within each cell were identified using mitochondrial masks, visible as confetti-like pseudo-coloring and string-like morphology, facilitating segmentation and characterization of each mitochondrion within each cell with unparalleled resolution (**Figure 4A-B**).

**Figure 4.**
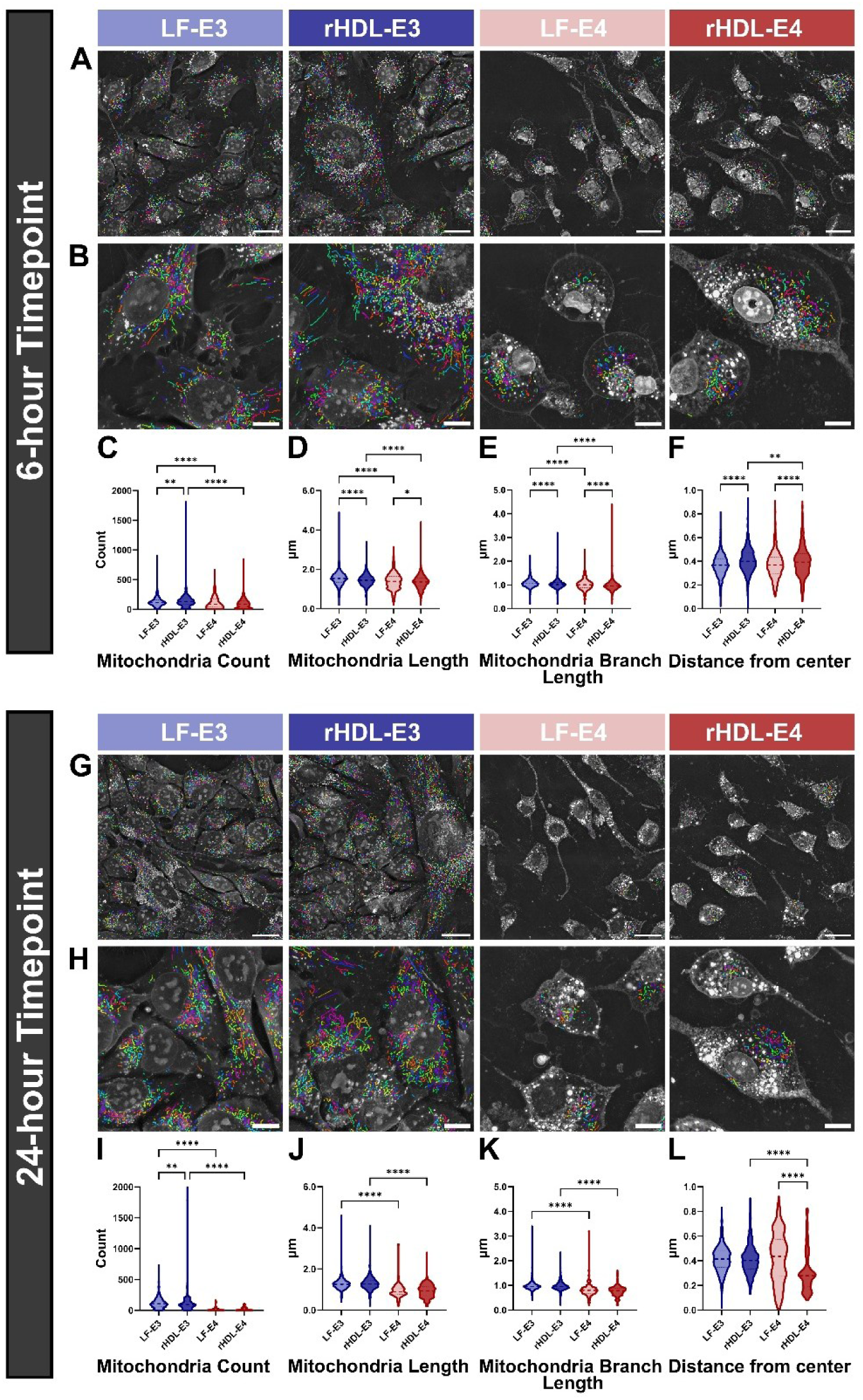
ApoE4 promotes mitochondrial fragmentation. Mitochondrial characterization of human microglia cells treated with lipid-free (LF-E3, LF-E4) or lipid-bound ApoE3 or ApoE4 (rHDL-E3, rHDL-E4) at 6 h (**A-F**) and 24 h (**G-L**). **A.** Mitochondrial masks at 6 h. Scale bars represent 25 µm. **B.** Zoomed-in mitochondrial masks at 6 h. Scale bars represent 10 µm. **C.** Mitochondria count per cell (6 h). **D.** Mitochondria length (6 h). **E.** Mitochondria branch length (6 h). **F.** Distance from cell center (6 h). **G.** Representative mitochondria masks at 24 h. Scale bars represent 25 µm. **H.** Zoomed-in mitochondrial mask at 24 h. Scale bars represent 10 µm. **I.** Mitochondria count per cell (24 h). **J.** Mitochondria length (24 h). **K.** Mitochondria branch length (24 h). **L.** Distance from cell center (24 h). Violin plots show median, upper, and lower quartiles (n**=**3, 50-100 cells). *\*p<0.01, **p<0.001, ****p<0.0001*.

By 6 h, mitochondria appear sparser with shorter ‘strings’ in cells treated with LF-E4 or rHDL-E4 compared to cells treated with LF-E3 or rHDL-E3 (**Figure 4A-B**). LF-E4 or rHDL-E4 supplementation also resulted in reduced mitochondrial count (**Figure 4C**) and length (**Figure 4D**) compared to cells treated with LF-E3 or rHDL-E3. In addition, cells treated with LF-E4 or rHDL-E4 exhibited lower mitochondrial branch length, indicative of mitochondrial fragmentation (**Figure 4E**). Taken together, these findings suggest that ApoE4 impairs mitochondrial dynamics, regardless of lipidation state. However, by 6 h, cells treated with rHDL-E3 or rHDL-E4 exhibited mitochondria that were further from the cell center (**Figure 4F**), highlighting a lipid-specific response to mitochondrial processing and subsequent localization within the cell.

By 24 h, the mitochondrial disruption seen at 6 h was more pronounced. Holotomography imaging revealed much sparser mitochondria following LF-E4 and rHDL-E4 supplementation (**Figure 4G-H**). Again, this was supported by quantification of mitochondrial dynamics, with LF-E4 and rHDL-E4 treatment reducing mitochondrial count (**Figure 4I**), length (**Figure 4J**), and branch length (**Figure 4K**), indicating mitochondrial fragmentation and thereby dysfunction in response to ApoE4, regardless of lipidation status. Overall, these imaging data suggest that treating HMg cells with ApoE4 leads to the accumulation of larger LDs, reflecting reduced lipid mobilization, which may be preceded by mitochondrial fragmentation and dysfunction.

### 3.6. Prolonged exposure to ApoE increases the abundance of proteins involved in lipid and cholesterol synthesis and interferon signaling

To probe the molecular mechanisms driving microglial responses to various ApoE treatments, we performed global proteomics analysis to quantify differences in protein abundance in the HMgs across the various ApoE treatment conditions (**Figure 5A**). Following peptide alignment, we identified 7,221 proteins across the experimental groups (**Supplementary Table 2**). A multivariate analysis using partial least squares–discriminant analysis (PLS-DA) revealed substantial overlap among the proteomic profiles of the different treatment groups (**Figure 5B**), but clear separation between the 6-hour and 24-hour timepoints. While several proteins exhibited changes in abundance across conditions at 6 hours (**Supplementary Table 2**), separation was minimal across groups indicating limited condition-specific differences in the proteomes. In contrast, PLS-DA suggested larger differences existed at the 24-hour timepoint. Indeed, PLS-DA of the 24-hour data revealed markedly enhanced separation between the treatment groups (**Figure 5C**), indicating the emergence of more distinct proteomic signatures with prolonged ApoE exposure. This temporal effect is consistent with our imaging data, which showed more pronounced alterations in cellular morphology, LD abundance, and mitochondrial dynamics following 24 h of ApoE exposure. Collectively, these results suggest that the primary driver of proteomic variance is the duration of ApoE supplementation rather than ApoE isoform or lipidation status.

**Figure 5.**
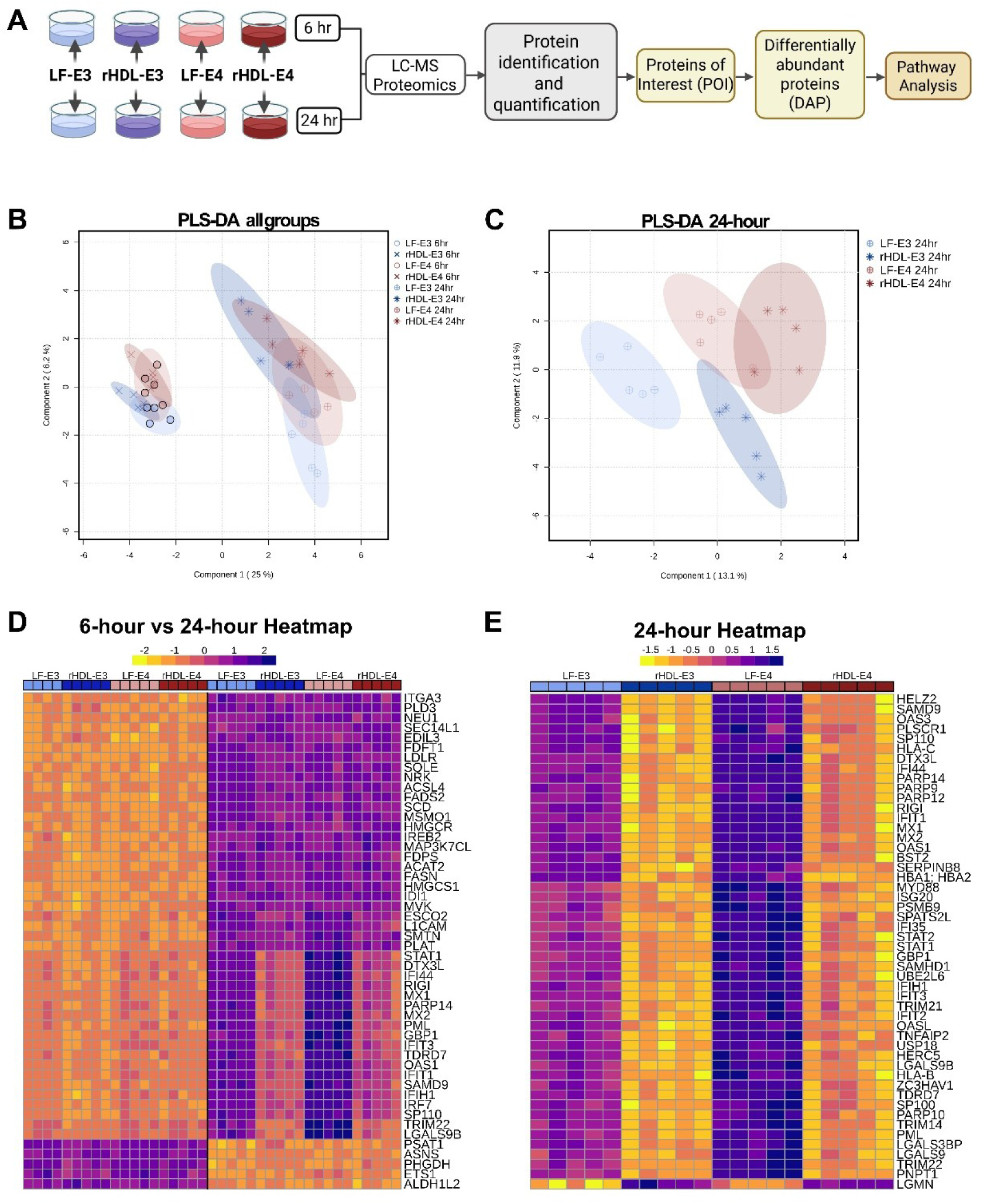
ApoE exposure remodels the microglial proteome by 24 hrs. **A.** Schematic of proteomic analysis of human microglia treated with lipid-free and lipid-bound ApoE3 or ApoE4 at 6 and 24 h. **B.** PLS-DA analysis of global proteomics across all ApoE treatment conditions and time points. **C.** PLS-DA analysis of global proteomics across all ApoE treatment conditions at the 24 h time point only. **D.** Heatmap of global proteomics across all ApoE treatment conditions showing the top 50 differentially abundant proteins. **E.** Heatmap of the top 50 differentially abundant proteins between the four ApoE treatment groups at the 24-h. Color gradients are associated with the normalized concentration (values closer to 2 have a higher concentration).

To define the proteomic signatures of these time-dependent differences, we visualized the top 50 differentially abundant proteins (DAPs) at both 6 and 24 h exposure to lipid-free and lipid-bound ApoE. We found statistically significant DAPs across the four treatment conditions. By 24 hours there was a robust increase in proteins involved in lipid and lipoprotein trafficking (LDLR, SEC14L1), fatty acid and triglyceride synthesis (FASN, SCD, ACSL4), and cholesterol biosynthesis and esterification (HMGCS1, ACAT2) (**Figure 5D**). The coordinated induction of lipid metabolic pathways is consistent with our imaging analyses and supports the notion that prolonged ApoE exposure leads to increased ApoE availability which promotes LD biogenesis and expansion in microglia. Extended ApoE treatment induced a pronounced inflammatory repolarization characterized by increased expression of interferon (IFN) signaling components (STAT1, MX2, IFITs, OAS1) and antiviral innate immune regulators (RIG-I, PARP14) (**Figure 5D**). Notably, these immune-associated proteins were most strongly elevated in cells supplemented with LF-E4, consistent with the idea that lipid-poor ApoE4 preferentially drives neuroinflammatory reprogramming.

While the majority of DAPs increased following 24 h ApoE exposure, a subset of proteins exhibited decreases in abundance. These included enzymes involved in amino acid biosynthesis and cytosolic NADPH production (PSAT1, PHGDH), transcriptional regulation of nuclear-encoded mitochondrial genes (ETS1), and mitochondrial NADPH generation (ALDH1L2). The coordinated reduction of these pathways suggests impaired mitochondrial metabolic support with increasing exposure to ApoE, a conclusion that aligns with our imaging-based observations of altered mitochondrial dynamics.

To further differentiate proteomic remodeling by LF-E3, LF-E4, rHDL-E3, and rHDL-E4, we compared the top 50 proteins with altered abundance after between the four treatment groups specifically at the 24 h exposure time point (**Figure 5E**). We observed proteins involved in antiviral (PARP14, PARP9, RIG1, MX1) and IFN signaling (STAT1, STAT2, SP100) were markedly increased in response to LF-E3 and LF-E4 but were most elevated in response to exposure to LF-E4.

To further determine whether increased lipid metabolism and interferon signaling were driven by extended ApoE exposure, we compared the proteomes of all ApoE-treated conditions to untreated control cells (**Supplementary** Fig. 2A). While minimal differences were observed at 6 h, by 24 h all groups, including controls, exhibited a time-dependent increase in proteins involved in lipid and lipoprotein metabolism (ACSL4, LDLR, FASN). This suggests that extended time in culture contributes to enhanced lipid processing, underscoring the importance of longitudinal metabolic assessment. In contrast, upregulation of immune and antiviral signaling proteins remained specifically associated with lipid-free ApoE treatments (**Supplementary Fig. 2A–B**). Consistent with this, the proteome of control cells most closely resembled that of rHDL-E3–treated cells, and to a lesser extent rHDL-E4–treated cells (**Supplementary Fig. 2A–D**). Pathway analysis further supported a robust association between lipid-free ApoE treatments and inflammatory signaling, (**Supplementary Fig. 2E–F**). Notably, LF-E4 uniquely induced PPAR signaling and fatty acid metabolism pathways, an effect not observed with LF-E3 (**Supplemental Figure 2F**). Together, these data indicate that the duration of ApoE exposure exerts a strong effect on microglial proteomic remodeling, which is most pronounced in response to lipid-free ApoE.

### 3.7. Lipid-free ApoE4 polarizes human microglia towards an antiviral, Type 1 interferon response

Given the marked impact of lipid-free ApoE on inflammatory polarization of human microglia, we next asked whether this immune response was stronger in response to LF-E4 treatment compared to LF-E3 treatment. PLS-DA demonstrated a clear isoform-dependent separation of proteomic signatures by 6 h (**Supplementary Fig. 3**) and 24 h exposure to LF-E3 versus LF-E4 treatment (**Figure 6A**). Examination of variable importance in projection (VIP) scores identified a discrete subset of proteins that contributed most strongly to this separation (**Figure 6B**). Moreover, differential abundance and hierarchical clustering analyses showed robust upregulation of IFN signaling components (STAT1, and IFIT1), antiviral proteins (MX1, OAS1-2, PARP14), and antigenic processing and presentation factors (HLA-B, PSMB8) in response to LF-E4 relative to LF-E3 treatment (**Figure 6C-D**).

**Figure 6.**
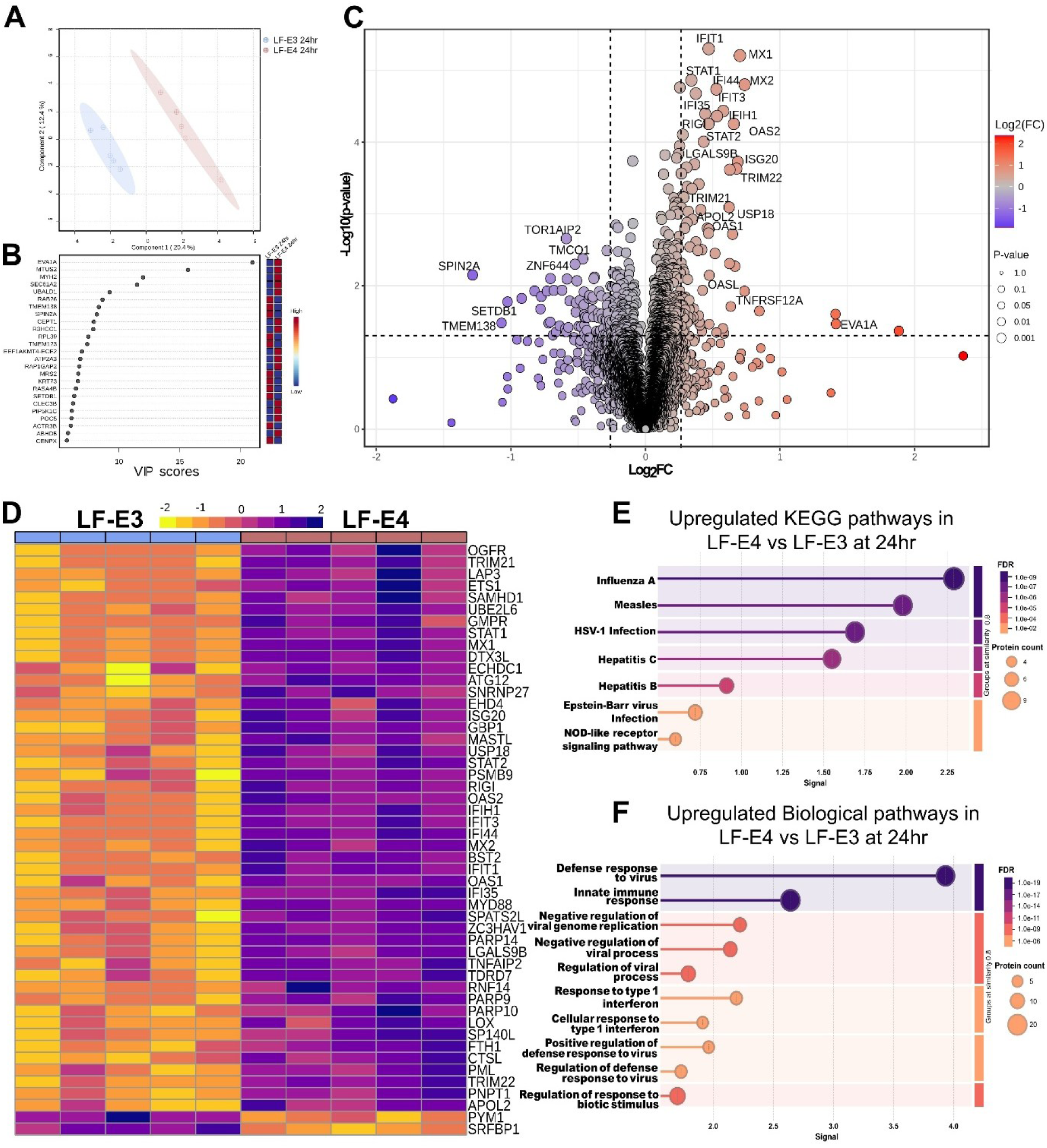
Lipid-free ApoE4 polarizes human microglia towards an antiviral, type 1 interferon response. **A.** PLS-DA plot of LF-E3 vs. LF-E4 treatment outcomes at the 24-h time point. **B.** VIP score table displaying the top 25 proteins driving differences in the PLS-DA plot. **C.** Volcano plot of LF-E3 vs. LF-E4 with x-axis representing log_2_FC and y-axis representing -log_10_(p-value Purple indicates downregulated proteins in response to LF-E4 treatment, while orange indicates upregulated proteins in response to LF-E4 treatment, at 24 h. **D.** Heatmap of the top 50 differentially abundant proteins in the 24 h group. The color gradient is associated with the normalized concentrations. **E.** Upregulated Kegg pathway analysis of DAPs in response to LF-E4 vs. LF-E3 treatment (upregulated in E4) at 24 h. **F.** Upregulated biological processes analysis of DAPs in response to LF-E4 vs. LF-E3 treatment (upregulated in E4) at 24 h.

Pathway enrichment analysis further confirmed significant activation of antiviral and innate immune pathways, with KEGG terms such as Influenza A, Herpes Simplex Virus-1 (HSV-1) infection, and NOD-like receptor (NLR) signaling prominently enriched in response to LF-E4 treatment (**Figure 6E**). ‘Biological processes’ pathway analyses revealed that LF-E4 elevated the abundance of proteins involved in innate immune defense, response to IFN, and regulation of viral processes (**Figure 6F**). To ensure robust pathway identification, pathway analysis was only performed on proteins that reached a stringent level of significance (p_adj_<0.1, **Supplementary Table 2**). The proteins driving these pathways are provided in **Supplementary Tables 3A-B.** Together, these data indicate that while the duration of ApoE exposure broadly governs microglial proteomic remodeling, LF-E4 acts as a potent and specific trigger of IFN and antiviral protein signatures.

### 3.8. Lipid-free ApoE3 drives antiviral and interferon signaling while suppressing proteostasis in human microglia

To determine how ApoE3 lipidation state influences human microglial responses, we compared proteomic signatures from cells treated with LF-E3 versus with rHDL-E3 over at 6 h (**Supplementary** Fig. 5) and 24 h. PLS-DA revealed clear separation between LF-E3 and rHDL-E3 treated HMg cells, indicating that ApoE3 lipidation state robustly shapes the microglial proteome by 24 h (**Figure 7A**). VIP scores identified a discrete subset of proteins driving this separation, including antiviral proteins previously described (OAS2, MX1, MX2) (**Figure 7B**). Differential expression analysis further supported these findings and demonstrated a pronounced skew toward proteins upregulated in LF-E3-treated cells, with relatively fewer proteins downregulated (**Figure 7C**).

**Figure 7.**
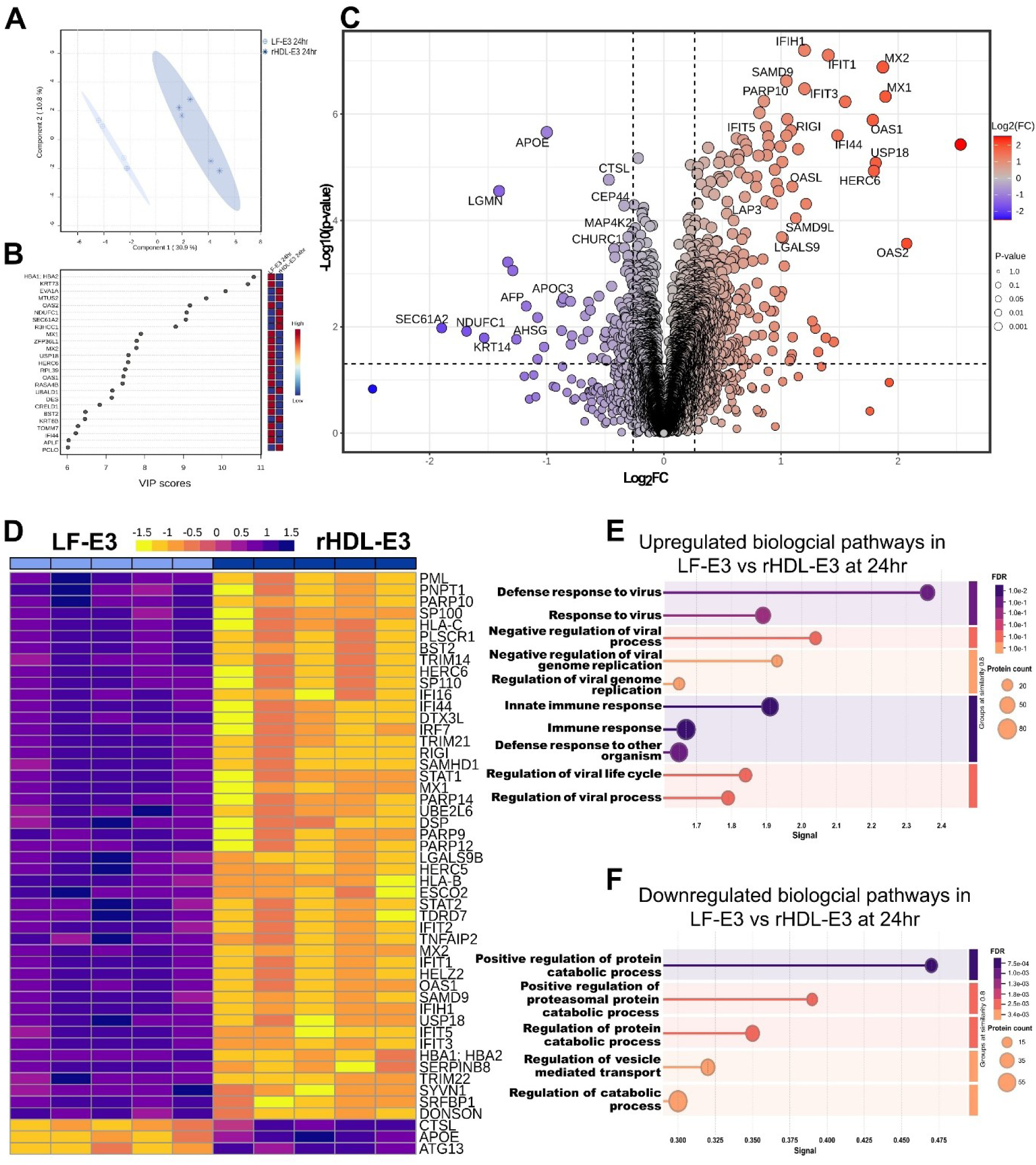
Lipid-free ApoE3 polarizes human microglia towards antiviral signaling and away from proteostasis. **A.** PLS-DA plot of LF-E3 vs. rHDL-E3 treatment outcomes at the 24-h time point. **B.** VIP score table displaying the top 25 proteins driving differences in the PLS-DA plot. **C.** Volcano plot displaying LF-E3 vs. rHDL-E3 with x-axis representing log_2_FC and y-axis representing -log_10_(p-value). Purple indicates downregulated proteins in response to LF-E3 and orange indicates upregulated proteins in response to LF-E3 treatment, at 24 h. **D.** Heatmap of the top 50 DAP proteins with color gradients associated with the normalized concentrations. **E.** Upregulated biological pathway analysis of DAPs in response to LF-E3 vs. rHDL-E3 treatment (upregulated in LF-E3) at 24 h. **F.** Downregulated biological processes analysis of DAs in response to LF-E3 vs. rHDL-E3 treatment (upregulated in LF-E3) at 24 h.

Consistent with these observations, hierarchical clustering of the top 50 DAPs revealed distinct patterns between the LF-E3 and rHDL-E3 treatment conditions (**Figure 7C**). LF-E3 treatment was characterized by coordinated increases in proteins involved in antiviral innate immunity and IFN signaling, including canonical IFN-stimulated genes and viral restriction factors (MX1, MX2, IFIT1–3, OAS1-2). In contrast, rHDL-E3–treated cells exhibited relatively higher abundance of proteins associated with lysosomal proteolysis and autophagy (CSTL, ATG13), suggesting a shift towards endolysosomal activity and stress-adaptive remodeling when ApoE3 is presented in a lipidated context.

Pathway enrichment analysis of proteins with increased abundance in LF-E3- versus rHDL-E3-treated microglia confirmed significant activation of antiviral and immune defense programs. Enriched biological pathways included defense response to virus, response to virus, negative regulation of viral replication, innate immune response, and IFN signaling (**Figure 7E, Supplementary Table 4A**). These data indicate that LF-E3 preferentially engages immune surveillance and antiviral programs in human microglia. Conversely, pathway enrichment analysis of proteins with decreased abundance, in response to LF-E3 relative to rHDL-E3 treatment, revealed significant suppression of biological processes related to protein metabolism and vesicular trafficking, including positive regulation of protein metabolism, catabolic processes, and regulation of protein turnover (**Figure 7F, Supplementary Table 4B**). Together, these findings suggest that LF-E3 drives a microglial state biased toward antiviral and IFN-mediated immune signaling, while concomitantly attenuating exocytic and endocytic pathways associated with membrane trafficking and proteostasis.

### 3.9. Lipid-free ApoE4 polarizes microglia to interferon and pattern recognition signaling and away from proteostasis programs

To assess how the lipidation state of ApoE4 alters microglial responses, we compared proteomic signatures in cells treated with LF-E4 or rHDL-E4 over the course of 6h (**Supplementary Fig. 6**) and 24 h (**Figure 8**). PLS-DA revealed a clear separation between treatment groups (**Supplementary Fig. 6A, Figure 8A**), driven by upregulation of proteins involved in IFN signaling, antiviral responses (IFIT1/2/3, MX1, OAS1) and fatty acid elongation (ELOV1) in response to 24 h LF-E4 exposure (**Figure 8B**). Differential abundance analyses corroborated this pattern, showing a pronounced shift toward proteins increased in response to LF-E4 exposed microglia treatment (**Figure 8C**), including key IFN-stimulated genes and pattern-recognition components (STAT1, IRF9, IFIH1/MDA5, DDX58/RIG-I, IFITM3, ISG15, HERC5, BST2). Proteins with increased abundance when expressed in response to exposed torHDL-E4 treatment mapped to lysosomal function and proteolysis (LGMN, SERPINC1), as well as resistance to apoptosis (TNFRSF10D).

**Figure 8.**
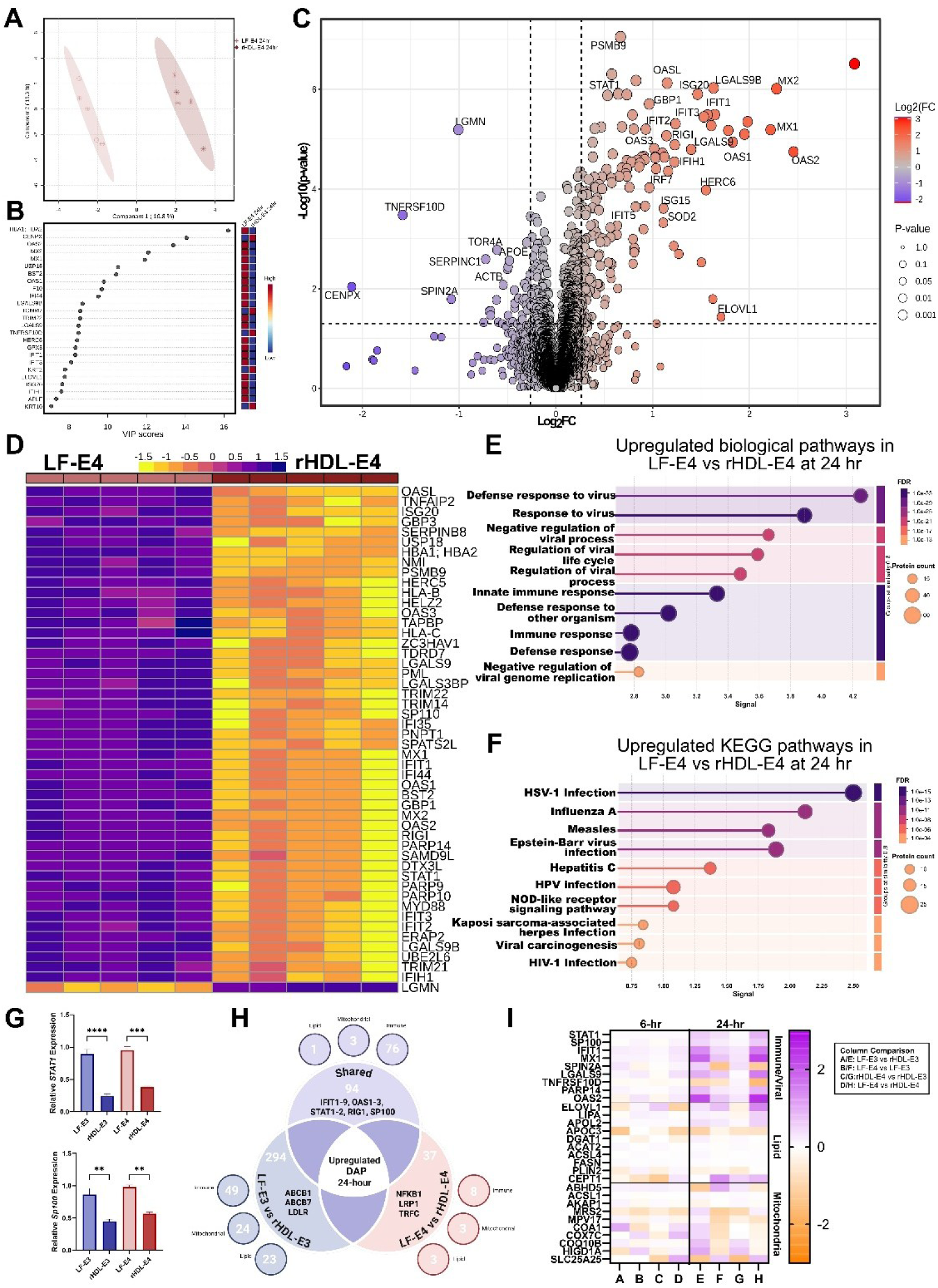
Lipid-free ApoE4 polarizes microglia to interferon and pattern recognition signaling and away from proteostasis programs. **A.** PLS-DA plot of LF-E4 vs. rHDL-E4 treatment at the 24-h time point. **B.** VIP score table displaying the top 25 proteins driving differences in the PLS-DA plot. **C.** Volcano plot displaying LF-E4 vs. rHDL-E4 with x-axis representing log_2_FC and y-axis representing -log_10_(p-value). Purple indicates downregulated proteins in response to LF-E4 treatment, while, orange indicates upregulated proteins in response to LF-E4 treatment, at 24 h. **D.** Heatmap of the top 50 DAP in the 24 h group. The color gradient is associated with the normalized concentration. **E.** Upregulated biological pathway analysis of DAPs in response to LF-E4 vs. rHDL-E4 treatment upregulated in LF-E4) at 24 h. **F.** Upregulated KEGG pathway analysis of DAPs in response to LF-E4 vs. rHDL-E4 treatment (upregulated in LF-E4) at 24 h. **G.** Relative gene expression of STAT1 and SP100. **H.** Venn diagram of overlapping DEPs between LF-E3 vs. rHDL-E3 treatment compared to LF-E4 vs. rHDL-E4 treatment. **I.** Heatmap of proteins of interest. Columns represent comparisons between groups at the 6- and 24 h time points. Rows indicate proteins categorized by function. Color scale indicates fold enrichment representing the left group versus the right group’s downregulation shown by a purple gradient and upregulation by an orange gradient. Data is mean ± SEM, n**=**(3) *, **p<0.001, ****p<0.0001*.

Comparing the top 50 DAPs revealed coordinated induction of antiviral modules under LF-E4 vs. rHDL-E4 treatment (**Figure 8D**), with elevated MX1/MX2, IFIT1–3, and OAS1/OAS2, while rHDL-E4 treatment favored proteolysis (LGMN). Pathway enrichment of DAPs increased in response to LF-E4 treatment highlighted defense response to virus, response to virus, negative regulation of viral replication, innate immune response, and IFN signaling (**Figure 8E, Supplementary Table 5A**), driven by STAT1, IRF9, DDX58, IFIH1, IFIT1–3, and OAS1/2. KEGG terms likewise emphasized viral infection pathways (HSV-1 infection, Influenza A, Measles, Hepatitis C/B, Epstein–Barr virus) and NLR signaling (**Figure 8F, Supplementary Table 5B**), reflecting shared pattern-recognition and IFN signatures (DDX58/RIG-I, IFIH1/MDA5, ISG15).

Orthogonal validation of gene expression confirmed the increased presence of IFN-related transcripts *Stat1* and *Sp100* in response to LF-E3 or LF-E4 treatment versus rHDL-E3 or rHDL-E4 treatment (**Figure 8G**), consistent with proteomic evidence of IFN pathway activation when ApoE is in its lipid-free state. Elevation of these antiviral transcripts was exacerbated in the presence of LF-E4 and rHDL-E4. Given the marked overlap in proteomic signatures in response to LF-E3 and LF-E4 treatment, we then compared POIs upregulated following treatment with LF-E4 versus rHDL-E4 to those upregulated following treatment with LF-E3 versus rHDL-E3 (**Figure 8H**). We found 94 proteins similarly upregulated in response to LF-E3 and LF-E4 treatment, with the majority of them involved in immune signaling (IFIT1-9, OAS1-3, STAT1-2, RIG1, SP100). We also identified 294 proteins uniquely upregulated in response to LF-E3 treatment, demonstrating a more diverse response, including 23 proteins involved in lipid signaling (LDLR, ABCB1) and 24 proteins involved in mitochondrial function. In contrast, there were only 37 proteins uniquely upregulated in response to LF-E4 treatment, including NFKB1 and LRP1 (**Figure 8H**). This finding indicates that while LF-E4 treatment polarizes microglia to an IFN dominant state, LF-E3 treatment induces a more diverse response, indicative of an attempt at reaching homeostasis and consistent with our imaging data showing higher cell health indices in microglia treated with LF-E3 and rHDL-E3.

Finally, while the global proteomic analysis revealed a robust induction of IFN signaling that largely dominated the response, this effect likely obscured more subtle changes in pathways that may underlie the increased LD accumulation and impaired mitochondrial dynamics observed in our live-cell imaging data. To address this, we reanalyzed our POIs and generated a temporal heat map to specifically examine changes in the abundance of proteins involved in lipid metabolism and mitochondrial function across ApoE treatment conditions. In addition to sustained IFN signaling, this analysis revealed upregulation of lipid processing enzymes (ELOVL1, LIPA) and concomitant downregulation of mitochondrial proteins involved in ion homeostasis and bioenergetic support (MRS2), in response to both LF-E4 and rHDL-E4 treatment. These proteomic shifts are consistent with our imaging results of enlarged LDs and increased mitochondrial fragmentation in cells under these conditions (**Figure 8I**). Together, these data indicate that lipid-free ApoE4 consistently polarizes human microglia toward an IFN-centered antiviral state, while lipid-free ApoE3 leads to a more heterogenous microglial response, which may facilitate cellular adaptation to mitigate cellular dysfunction.

## 4. Discussion

Although *APOE ε4* is the strongest common genetic risk factor for AD, the mechanisms linking ApoE4 to AD neuropathogenesis remain incompletely defined. ApoE4 has been implicated in impaired lipid transport and microglial dysfunction, characterized by lipid droplet (LD) accumulation and reduced phagocytosis^22,52^. However, prior studies have focused on endogenous ApoE4 expression, overlooking the impact of ApoE lipidation on microglial metabolism and function. To address this gap, we compared the impact of lipid-free (LF) and lipid-bound (rHDL) ApoE3 and ApoE4 on human microglia, combining live-cell holotomography with unbiased proteomics. Across conditions, ApoE4 reduced cell health, promoted fewer but larger LDs—especially when lipidated—and induced mitochondrial fragmentation. Proteomic profiling demonstrated induction of lipid biosynthesis following prolonged ApoE exposure, alongside a strong IFN response elicited by LF-ApoE, and most prominently LF-ApoE4. Together, these data indicate that ApoE4’s lipidation state is a critical determinant of microglial immunometabolic programming, with rHDL-ApoE4 driving metabolic dysfunction, and LF-ApoE4 amplifying inflammatory signaling.

Here, we supplemented murine and human microglia with LF-ApoE3, LF-ApoE4, rHDL-ApoE3, or rHDL-ApoE4. While previous studies have utilized relevant models, such as human iPSC-derived microglia or murine models of ApoE4 expression, it is challenging to empirically determine the effects of either LF or rHDL-ApoE in these studies due to the heterogeneous nature of endogenously expressed ApoE-containing particles. In addition, particle characterization is rarely considered, especially *in vivo*, due to the technical challenges of isolating sufficient quantities to effectively assess lipid and protein content. To overcome this scientific roadblock, we generated either ApoE3 or ApoE4-containing HDL-like particles with precisely controlled lipid and protein composition. These homogenous particles contain two ApoE proteins, consistent with the current model of ApoE lipidation, in which two ApoE molecules form antiparallel α-helical chains around lipoprotein discs, adopting a “double-belt” structure^54^. This structure differs considerably from that of LF-ApoE, which exhibits a compact, amphiphilic, α-helical structure, resulting in reduced surface area, less accessible lipid- and receptor-binding regions, and a propensity to aggregate^55^. While the structural characteristics of LF-ApoE4 are thought to promote Aβ oligomerization, plaque deposition, and neurotoxicity in AD^56^, how ApoE isoform and lipidation status directly affect microglia has, to date, remained elusive.

Here, we show that rHDL-ApoE4 can enter the cell more readily than rHDL-ApoE3. This is perhaps surprising, given the accepted dogma that ApoE4 is less efficient at transporting cholesterol than ApoE2 or ApoE3^57^. However, this data is consistent with recent reports highlighting a stronger interaction between ApoE4 and lipoprotein receptors, such as triggering receptor on myeloid cells 2 (TREM2)^58^ and low-density lipoprotein receptor (LDLR)^59^. In addition, recent studies have shown that decreased binding between LDLR and rHDL-ApoE2 or the protective Christchurch variant (which occurs on an ApoE3 background) reduces the uptake of pathogenic lipid cargos (e.g., cholesteryl ester [CE]) and prevents cellular dysfunction characterized by lipid peroxidation and disordered lysosomal processing^60^. Therefore, it is reasonable to suggest that a strong, energetically favorable interaction between rHDL-ApoE4 and lipoprotein receptors may increase lipid delivery and intracellular sequestration of ApoE4, contributing to microglial reprogramming in AD pathology. Notably, our rHDL particles contain equivalent phospholipid and ApoE content (with no cholesterol or CE), suggesting that the observed preferential uptake of ApoE4 reflects optimized lipid handling that becomes apparent only when ApoE4 is fully lipidated.

The ability to determine how ApoE affects the metabolic reprogramming of microglia has historically been challenging due to technical limitations in longitudinal imaging of lipids in live cells and in monitoring lipid processing *in situ* in real time. To overcome this, we used label-free live-cell holotomography, which uses the change in refractive index as light passes through a given cell to construct extremely high-resolution images of cellular and intracellular structures^47–49^. In the present study, we used this technique to image and quantify cellular morphology, LD accumulation, and mitochondrial dynamics in real time as cells responded to LF- and rHDL-ApoE3 or ApoE4. Morphological analysis revealed that cells treated with ApoE3 retained typical morphology even after 24 h of treatment, regardless of lipidation state. In contrast, cells treated with ApoE4 shifted from a larger, more ramified state to a smaller, more circular amoeboid morphology over the 24-h period, an effect that was pronounced when cells were treated with rHDL-ApoE4. Importantly, rHDL-ApoE3 and rHDL-ApoE4 were compositionally matched (200 μM POPC, 0.01mg/mL ApoE), indicating that the observed morphological changes truly reflect lipidation-state and ApoE-isoform-specific effects, rather than disparities in particle dose. These findings are consistent with the notion that lipid delivery is enhanced when ApoE4 is fully lipidated, which may not always be in the case in neuroinflammatory and neuropathogenic disease.

Robust morphological alterations in ApoE-treated cells suggest that, once internalized, ApoE4 triggers an intracellular signaling cascade leading4 to cellular reorganization. This is consistent with recent studies in iPSC-derived endothelial cells showing that ApoE4 can impair endosomal processing and intracellular transport^61^. Endosomal and lysosomal dysfunction has also been reported in ApoE4-expressing microglia in rodent models of AD and in human AD brains. Notably, ApoE4 aggregates within lysosomes, which contributes to Aβ accumulation in immortalized, primary, and iPSC-derived microglia *in vitro*^61^. Aβ aggregation appears to be exacerbated in cells treated with ApoE bound to POPC and cholesterol, and even more so in response to lipidated ApoE4^62^. Our findings support these recent studies and the emerging hypothesis that while rHDL-ApoE4 may enter microglia more readily, once internalized rHDL-ApoE4 readily undergoes de-lipidation and aggregation, contributing to AD pathology.

In addition to high-resolution analysis of cellular morphology, the holotomography platform used in this study also facilitates the real-time quantification of LD dynamics^63–65^. Interestingly, we found that cells treated with ApoE3 had a slightly higher LD count than those treated with ApoE4, consistent with recent studies showing that ApoE3-expressing human microglia have more LDs than microglia expressing ApoE4^22^. In contrast, our high-resolution analysis of LD dynamics showed that while cells treated with ApoE4 did not accumulate more LDs, they did, in fact, accumulate larger LDs. While LD expansion has not been previously demonstrated in microglia, an ApoE4-induced increase in LD size has recently been shown in human astrocytes^66^. Since fewer but larger LDs are characteristic of deficits in lipolysis^67–69^, our data suggests that once internalized, ApoE4—and perhaps more so rHDL-ApoE4—impairs the function of intracellular lipases, reducing lipid mobilization. This is further supported by our observation that cells treated with rHDL-ApoE4 have LDs that are further from the cell nucleus (more peripheral), highlighting impaired lipid turnover and spatial organization of lipid-processing machinery^69,70^. Given the recent studies described above, it is likely that this is due to ApoE4 aggregation within intracellular vesicles. Indeed, LDs within cells treated with rHDL-ApoE4 are described as more ‘tightly packed’ a marker of protein density indicative of protein aggregation and reduced lipid surface area, which inhibits lipase access and lipid mobilization^68^. Since ApoE4-mediated LD accumulation is known to inhibit microglial function and neuronal support^52^, our work further supports the notion that ApoE4 leads to the accumulation of large LDs, indicative of immunometabolic dysfunction that contributes to neurodegenerative disease pathology.

A key component of metabolic remodeling in microglia is the shift towards glycolysis and away from mitochondrial metabolism, which not only contributes to inflammatory polarization but also contributes to LD accumulation through impaired lipid catabolism and the generation of glycolytic intermediates that promote lipogenesis^71^. Another advantage of live-cell label-free holotomography is the ability to visualize and quantify mitochondrial number, morphology, and fragmentation, providing valuable insights into mitochondrial dynamics that are essential for interpreting the ApoE4-mediated effects on cell morphology and LD accumulation. Using this platform, we observed higher mitochondrial counts, lengths, and branch lengths in human microglial cells treated with LF- and rHDL-ApoE3. Since these mitochondrial characteristics are associated with a fusion-dominant morphology^72,73^, it is reasonable to suggest that mitochondria within ApoE3-treated cells are more functional. In contrast, microglia treated with LF- and rHDL-ApoE4 exhibited reduced mitochondrial number and branch length, indicative of mitochondrial fragmentation^73^. In addition, mitochondria clustered towards the nucleus in cells treated with rHDL-ApoE4. Since increased proximity to the nucleus is a hallmark of mitochondrial stress^74^, our findings indicate that lipid-bound ApoE4 induces mitochondrial stress and mitochondrial dysfunction. While the mechanisms underlying ApoE4-mediated mitochondrial fragmentation in microglia remain elusive, impaired mitochondrial dynamics have repeatedly been observed in astrocytes and neurons^75^, which is thought to involve ApoE4-mediated reorganization of contact sites between mitochondria and the endoplasmic reticulum^76,77^. Such cellular reorganization aligns with our observations showing that ApoE4 impacts mitochondrial dynamics and LD position more so than ApoE3.

While holotomography offers unique insights into LD and mitochondrial dynamics, high-resolution imaging studies cannot identify the cellular and molecular mechanisms underlying the lipidation and isoform-specific effects of ApoE on microglia. Therefore, to provide mechanistic insights, we performed global proteomics on microglia treated with LF- and rHDL-ApoE3 and ApoE4. Notably, with prolonged exposure to both isoforms of ApoE—regardless of lipidation status—we observed increased abundance of proteins involved in lipid and cholesterol biosynthesis, including fatty acid synthase (FASN) and acyl-CoA: cholesterol-acyltransferase (ACAT). Notably, FASN, the rate limiting enzyme in de novo lipogenesis, is increased in microglia from murine models of AD (5xFAD), and in the brains of individuals with AD^78,79^. This is consistent with the notion that AD brains show higher triglyceride levels, potentially due to microglial lipid accumulation^16^. Moreover, recent reports have also shown that pharmacological FASN inhibition can restore microglial lipids to control levels, supporting its therapeutic potential^79^. Similarly, pharmacological ACAT inhibition has been shown to prevent LD accumulation in 5xFAD mice expressing ApoE4, although it does not improve ApoE lipidation. Taken together, these results indicate that ApoE abundance, independent of isoform or lipidation state, can drive lipid biosynthesis in microglia, with effects likely exacerbated in neuroinflammatory contexts, such as in the presence of ApoE4^80^.

Perhaps the most striking observation from our proteomics analysis was the elevated expression of proteins involved in IFN and antiviral signaling (e.g., STAT1, SP100, IFIT, PARP14, and OAS2) following prolonged exposure to LF-ApoE. Detailed comparison between LF-ApoE3 and rHDL-ApoE3 versus LF-ApoE4 and rHDL-ApoE4 revealed that LF-ApoE had the strongest impact on inflammatory polarization of human microglia, regardless of isoform. However, when LF-ApoE3 and LF-ApoE4 were directly compared, we found that LF-ApoE4 resulted in an exacerbated IFN and antiviral response. Interestingly, the response to LF-ApoE3 appeared to be more heterogenous, implying an attempt at achieving cellular homeostasis. Importantly, the ApoE proteins in the LF and rHDL particles were identical, indicating that LF-ApoE4 particles alone were responsible for inflammatory remodeling. These data are consistent with the notion that ApoE4 may be relatively lipid-poor compared to ApoE2 or ApoE3 and may explain why ApoE4 is typically associated with greater inflammatory signaling in cells, mice, and humans^81–83^, leading to enhanced cellular vulnerability to neurodegeneration. In support of this notion, recent studies have shown that IFN signaling is elevated in primary ApoE4-expressing microglia^84^, and in microglia from multiple AD models^85^, highlighting the therapeutic potential of targeting this pathway. Indeed, the tendency of LF-ApoE to promote inflammatory polarization may be mechanistically distinct from the propensity of rHDL-ApoE4 to induce metabolic reprogramming and enhance lipid availability. Regardless, while ‘optimally-lipidated’ ApoE4 appears to enter cells more readily, and LF-ApoE4 is a more potent inflammatory signal, both states may be necessary, but not sufficient, for AD onset and progression.

Despite the strengths of our methodological approach, several limitations warrant consideration. Firstly, although human fetal microglial (HMG) cell lines are widely used and well suited for exploratory studies, they are immortalized and thus do not fully recapitulate the cellular complexity or physiological state of primary or patient-derived microglia. Nevertheless, robust global proteomic profiling in our workflow required approximately ∼500,000 cells per replicate (n = 5) per condition—a scale that exceeds the practical yield achievable from primary human or iPSC-derived microglia—necessitating the use of an immortalized human cell line. Importantly, several key protein pathways identified in our analysis have also been reported to be elevated in primary microglia, as well as microglia from murine models of AD and in human studies following ApoE4 exposure and expression, thereby supporting the biological relevance and validity of our experimental pipeline. Even so, future studies should aim to validate these pathways in primary and iPSC-derived microglial models to further strengthen translational relevance. Secondly, while holotomography is particularly advantageous for longitudinal observation of LDs in microglia, this technology has only previously been used to understand LD accumulation in cancer and metabolic disease, and therefore not been extensively applied to neurodegenerative disease models^86,87^. Nonetheless, holotomography holds significant promise as a gold-standard approach for assessing LDs and mitochondrial dynamics in live microglia. Finally, we appreciate that our rHDL particles do not contain cholesterol or CE, and contain only PLs and ApoE, therefore not perfectly recapitulating the lipid and protein composition of brain-derived lipoproteins. Despite these limitations, our findings lay critical groundwork for future studies examining how ApoE lipidation state and isoform contribute to microglial dysfunction in AD.

In summary, ApoE4 disrupts microglial homeostasis through distinct mechanisms determined by its lipidation state. Specifically, lipid-bound ApoE4 may drive metabolic dysfunction and elevated lipid supply, when fully lipidated, whereas lipid-free ApoE4 amplifies IFN and antiviral signaling. These findings underscore the critical need to account for lipidation status when investigating ApoE4-mediated pathology in AD and suggest that both lipid-bound and lipid-free states may contribute to AD neuropathogenesis.

## DATA AVAILABILITY

The data supporting the findings of this study are available upon reasonable request from the corresponding author K.D.B. The mass spectrometry proteomics has been deposited to the ProteomeXchange https://massive.ucsd.edu/ProteoSAFe/dataset.jsp?task=df6988282d714c6e8894927f09506ca3

## ACKNOWLEDGEMENTS

We would like to thank the personnel from the University of Colorado Anschutz Medical Campus Advanced Light Microscopy Core for imaging assistance. Imaging was performed in the Advanced Light Microscopy Core facility of the NeuroTechnology Center at the University of Colorado Anschutz Medical Campus, which is supported in part by Rocky Mountain Neurological Disorders Core Grant (P30NS048154) and by Diabetes Research Center Grant (P30 DK116073).

## Contributions

Conceptualization: K.D.B., J.T.M,

Experimentation/Investigation: T.G.S, S.S, K.M.B, R.W, H.B, V.P, C.O, A.S, K.A

Writing original draft: T.G.S, K.M.B, K.D.B, J.T.M

Writing-review & editing: T.G.S, K.D.B, J.E.H., R.H.E., K.G.S, C.S.N.

Supervision: K.D.B., J.T.M

