## Supplemental Figures and Tables for "ApoE Lipidation State Directs Immunometabolic Reprogramming of Human Microglia"

**Supplementary Table 1.** Nanolive Metric and Interpretation

| Nanolive Metric Label | Nanolive Interpretation | Our Metric Label | Our Interpretation in relation to microglia |
|---|---|---|---|
| Health Index | Numbers closer to 0 represent stressed cells; numbers closer to 1 represent healthy cells. | Cell Health Index | How healthy the cell is. |
| Area | Area in a single object. | Cell Area or LD Area | How big the microglia or the LD is. |
| Form Factor | Closer to 1 indicates a circular object. Value closer to 0 means elongated spaghetti like shapes. Calculated as: $\frac{4\pi * Area}{Perimeter^2}$ | Circularity | Morphological Parameter measuring how close of a circle the cell is. For microglia the more circular the more amoeboid. |
| Compactness | Objects that have multiple protrusions in different directions will have values greater than 1. Calculated as: $\frac{Perimeter^2}{4\pi * Area}$ | Ramification | High ramification can indicate a more homeostatic microglia vs low ramification which means more compact activated microglia |
| Mean Radial Distance Index | Describes how close an object is to the parent's cell center. Values closer to 0 indicate closer to the cell center, values closer to 1 indicate object is closer to the cell edge. | Distance from center | How close the object is to the cell center aka nucleus. |
| Mean Dry Mass Density | Dry mass (content of the object excluding water i.e. lipids) divided by the estimated object volume. | LD Packing | How tightly packed the LD is. A higher dry mass density means more lipids are packed into the same volume. |
| Count | Total number of objects in the parent cell . The object is in reference to the specific metric we are looking at such as lipid droplet or mitochondria | LD Count Mitochondria Count | Number of objects (LD or mitochondria) in the parent cell. |
| Mean Total Length | Mean length of mitochondria in the parent cell | Mitochondria Length | How long the average mitochondria is |
| Mean Branch Length | Average of the mean branch length per mitochondrion in the parent cell | Mitochondria Branch Length | How long the average branch of a mitochondrion is |

**Supplementary Table 2.** Protein counts

| 7221 Total Proteins Identified | | |
|---|---|---|
| **Comparison** | **p-value ≤0.05 (Proteins of Interest [POI])** | **p-adj ≤0.1 (Differentially abundant proteins [DAP])** |
| LF-E4 vs LF-E3 6hr | 1008 | 2 DAP<br>1 upregulated<br>1 downregulated |
| rHDL-E4 vs rHDL-E3 6hr | 680 | 0 |
| LF-E3 vs rHDL-E3 6hr | 918 | 0 |
| LF-E4 vs rHDL-E4 6hr | 1413 | 17 DAP<br>13 upregulated<br>4 downregulated |
| LF-E4 vs LF-E3 24hr | 639 | 27 DAP<br>27 upregulated |
| rHDL-E4 vs rHDL-E3 24hr | 326 | 0 |
| LF-E3 vs rHDL-E3 24hr | 1977 | 730 DAP<br>387 upregulated<br>343 downregulated |
| LF-E4 vs rHDL-E4 24hr | 776 | 155 DAP<br>130 upregulated<br>25 downregulated |

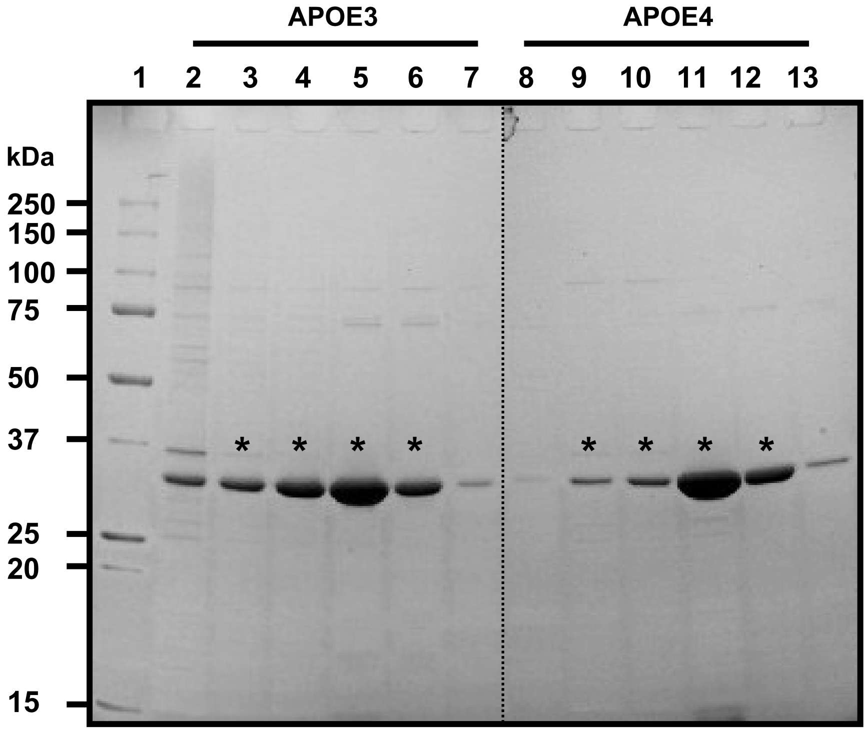


**Supplemental Figure 1. APOE protein purity.** Lyophilized APOE was unfolded in 3M guanidine-HCl, refolded by dialysis, and purified by size exclusion chromatography using a HiLoad 16/600 Superdex 200 pg column. 1 mL fractions were collected and 15 µl of each fraction were analyzed on an 10-20 % gradient SDS-PAGE under reducing conditions. Lanes 2-7 contain fractions from APOE3, while lanes 8-13 contain fractions from APOE4. Protein bands were visualized using Coomassie Blue staining. Fractions marked with asterisks indicate those that were pooled and concentrated for subsequent experimental use.

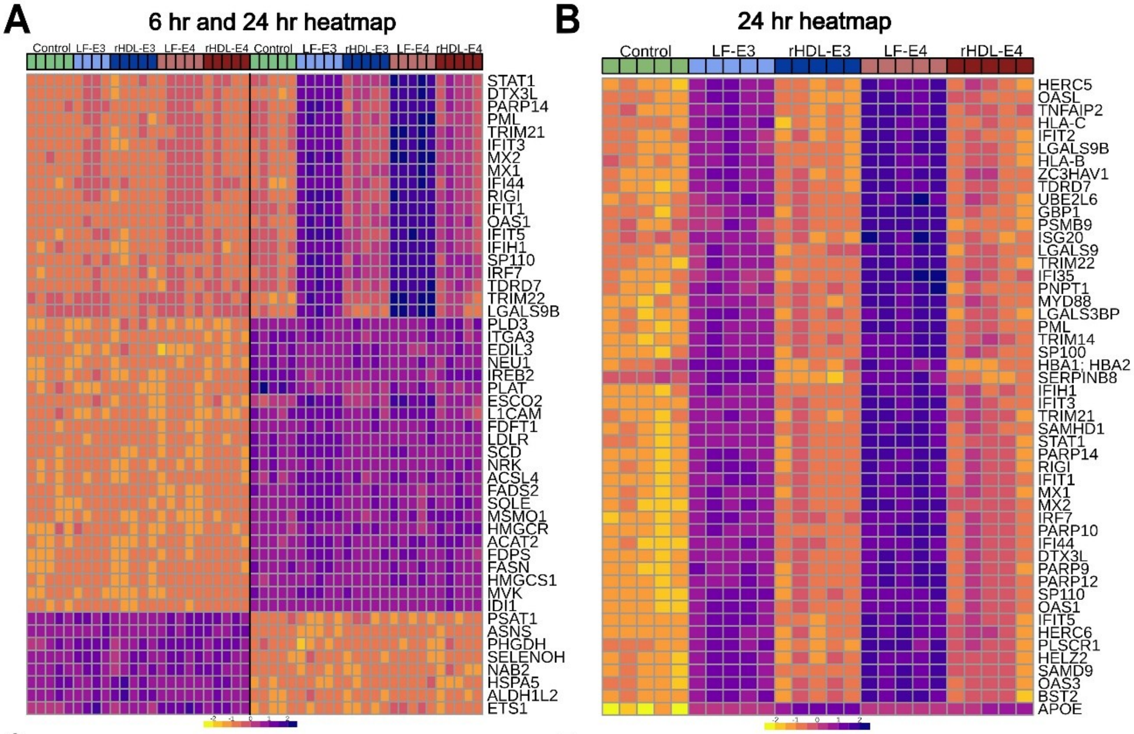


**C**

| Comparison | Proteins with p-value ≤0.05 (POI) | Proteins with p-adj ≤0.1 (DAP) |
|---|---|---|
| LF-E3 vs Ctrl 6hr | 720 | **1 DAP**<br>1 downregulated |
| rHDL-E3 vs Ctrl 6hr | 281 | 0 |
| LF-E4 vs Ctrl 6hr | 1406 | **274 DAP**<br>194 upregulated<br>81 downregulated |
| rHDL-E4 vs Ctrl 6hr | 597 | 0 |
| LF-E3 vs Ctrl 24hr | 1886 | **817 DAP**<br>383 upregulated<br>434 downregulated |
| rHDL-E3 vs Ctrl 24hr | 490 | **6 DAP**<br>2 upregulated<br>4 downregulated |
| LF-E4 vs Ctrl 24hr | 1083 | **272 DAP**<br>193 upregulated<br>78 downregulated |
| rHDL-E4 vs Ctrl 24hr | 660 | 0 |

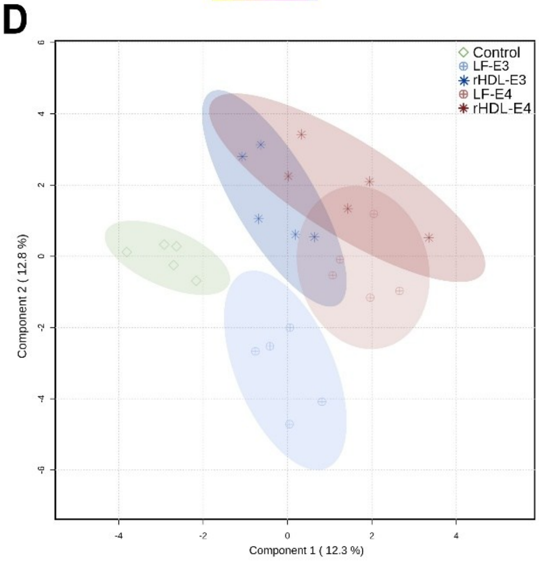


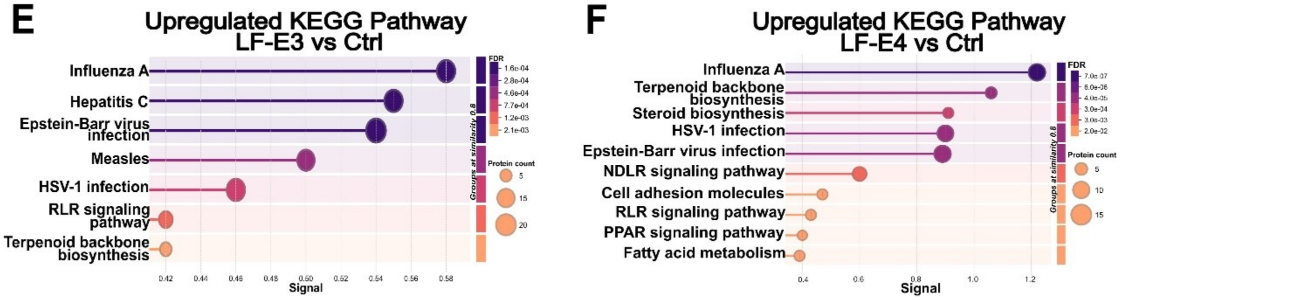


**Supplemental Figure 2. ApoE remodels the microglial proteome by 24 hours A.** Heatmap of global proteomics across all ApoE treatment conditions showing the top 50 differentially abundant proteins. **B**. Heatmap of the top 50 differentially abundant proteins between the four ApoE treatment groups and the control at the 24-h. Color gradients are associated with normalized concentration (values closer to 2 have a higher concentration) **C.** Table displaying the protein counts identified within each potential comparison. **D.** PLS-DA analysis of global proteomics across conditions and time points. **E.** Upregulated KEGG pathway analysis of DAPs in response to LF-E3 vs. Control (upregulated in LF-E4) at 24-h. **F.** Upregulated KEGG pathway analysis of DAPs in response to LF-E4 vs. control (upregulated in LF-E4) at 24 h.

**Supplementary Table 3A.** Upregulated KEGG pathways in LF-E4 vs LF-E3 at 24 h

| KEGG Pathway | Protein count | Background protein count | Strength | Signal | FDR | Proteins |
|---|---|---|---|---|---|---|
| Influenza A | 9 | 79 | 1.48 | 0.0155 | 5.46e-09 | PML,STAT2,MX2,OAS2,STAT1,DDX58,MX1,IFIH1,MYD88 |
| Measles | 8 | 78 | 1.43 | 0.0155 | 9.50e-08 | STAT2,MX2,OAS2,STAT1,DDX58,MX1,IFIH1,MYD88 |
| Herpes simplex virus 1 infection | 8 | 102 | 1.32 | 0.0155 | 4.62e-07 | BST2,PML,STAT2,OAS2,STAT1,DDX58,IFIH1,MYD88 |
| Hepatitis C | 7 | 87 | 1.33 | 0.0155 | 3.18e-06 | STAT2,MX2,OAS2,STAT1,IFIT1,DDX58,MX1 |
| Hepatitis B | 5 | 85 | 1.19 | 0.0155 | 0.0012 | STAT2,STAT1,DDX58,IFIH1,MYD88 |
| Epstein-Barr virus infection | 5 | 115 | 1.06 | 0.0155 | 0.004 | STAT2,OAS2,STAT1,DDX58,MYD88 |
| NOD-like receptor signaling pathway | 4 | 86 | 1.09 | 0.0155 | 0.0155 | STAT2,OAS2,STAT1,MYD88 |

**Supplementary Table 3B**. Upregulated Biological pathways in LF-E4 vs LF-E3 at 24 h

| Biological Pathway | protein count | background protein count | Strength | Signal | FDR | Proteins |
|---|---|---|---|---|---|---|
| Defense response to virus | 17 | 119 | 1.58 | 4.06 | 1.35e-19 | ZC3HAV1,BST2,PML,DTX3L,ISG20,STAT2,MX2,OAS2,PARP9,STAT1,IFIT1,IFIT3,DDX58,TRIM22,MX1,IFIH1,MYD88 |
| Innate immune response | 19 | 250 | 1.3 | 2.77 | 1.81e-18 | ZC3HAV1,BST2,PML,DTX3L,ISG20,STAT2,MX2,OAS2,PARP9,STAT1,IFIT1,IFIT3,DDX58,TRIM22,MX1,IFI35,PARP14,IFIH1,MYD88 |
| Regulation of response to biotic stimulus | 11 | 162 | 1.25 | 1.83 | 7.59e-09 | USP18,ZC3HAV1,PML,DTX3L,STAT2,PARP9,STAT1,IFIT1,DDX58,IFI35,PARP14 |
| Negative regulation of viral process | 8 | 57 | 1.57 | 2.27 | 3.65e-08 | ZC3HAV1,BST2,ISG20,OAS2,STAT1,IFIT1,MX1,IFIH1 |
| Regulation of viral process | 9 | 100 | 1.38 | 1.92 | 7.00e-08 | ZC3HAV1,BST2,ISG20,OAS2,STAT1,IFIT1,TRIM22,MX1,IFIH1 |
| Negative regulation of viral genome replication | 7 | 37 | 1.7 | 2.35 | 9.13e-08 | ZC3HAV1,BST2,ISG20,OAS2,IFIT1,MX1,IFIH1 |
| Response to type I interferon | 6 | 23 | 1.84 | 2.32 | 3.39e-07 | STAT2,OAS2,STAT1,IFIT1,MX1,MYD88 |
| Positive regulation of defense response to virus by host | 5 | 15 | 1.94 | 2.09 | 3.05e-06 | PML,DTX3L,PARP9,STAT1,DDX58 |
| Regulation of defense response to virus | 6 | 37 | 1.63 | 1.86 | 3.36e-06 | PML,DTX3L,PARP9,STAT1,IFIT1,DDX58 |
| Cellular response to type I interferon | 5 | 16 | 1.92 | 2.04 | 3.69e-06 | STAT2,OAS2,STAT1,IFIT1,MYD88 |

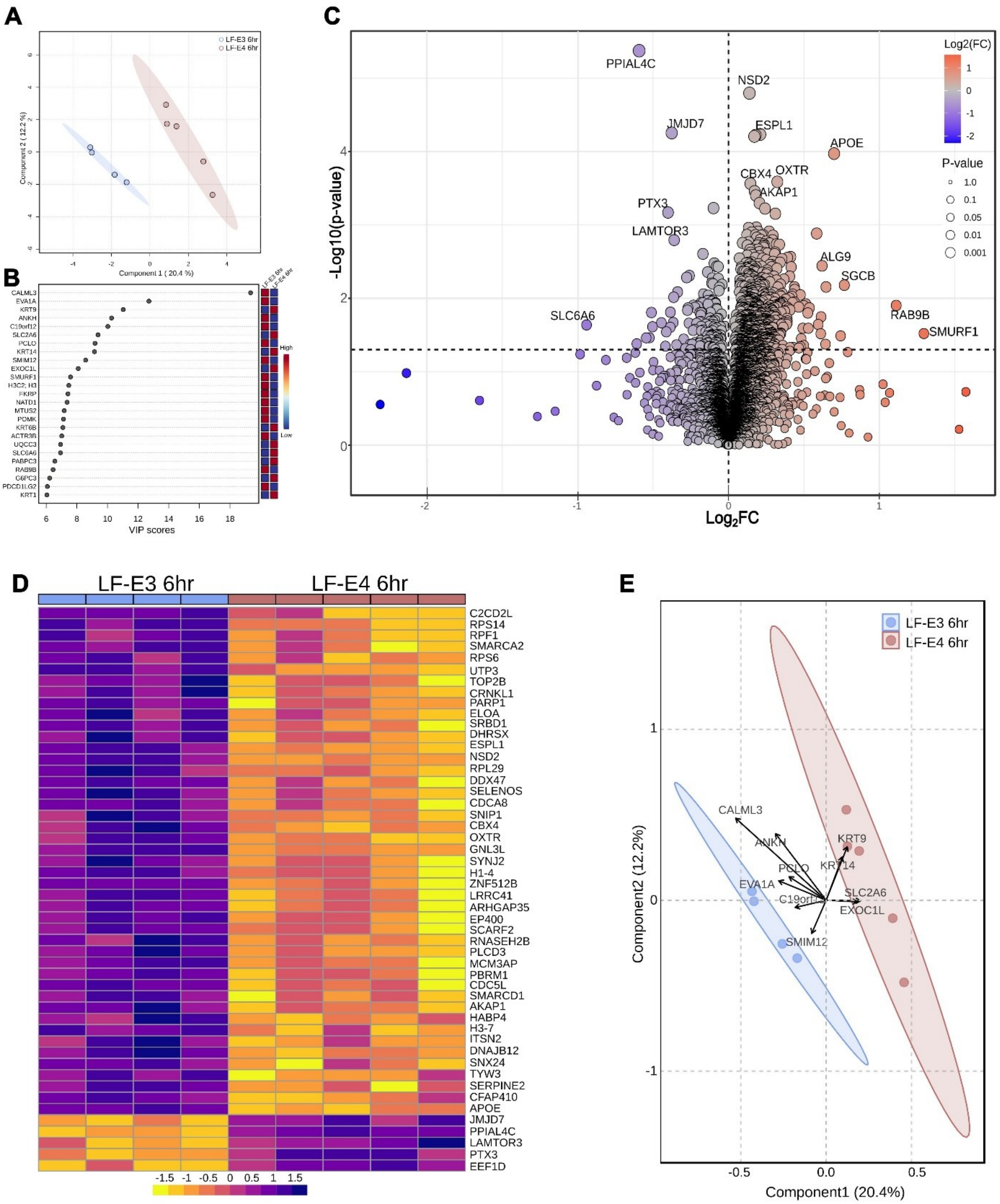


**Supplemental Figure 3. Comparative proteomics of microglia treated with either LF-E3 or LF-E4 for 6 hours. A**. PLS-DA plot of LF-E3 vs LF-E4 treatment outcomes at the 6 h time point. **B.** VIP score table displaying the top 25 proteins driving differences in the PLS-DA plot. **C.** Volcano plot displaying LF-E3 vs LF-E4 with x-axis representing $log_2$FC and y-axis representing $-log_{10}$(p-value). Purple indicates downregulated proteins in response to LF-E3 and orange indicates upregulated proteins in response to LF-E3 treatment, at 6 h. **D.** Heatmap of the top 50 DAP proteins with color gradients associated with the normalized concentrations. **E.** PLS-DA biplot with arrows representing gene loadings; direction and length indicate the contribution of specific proteins to the sample separation. Proteins pointing to a specific cluster are positively associated with that group.

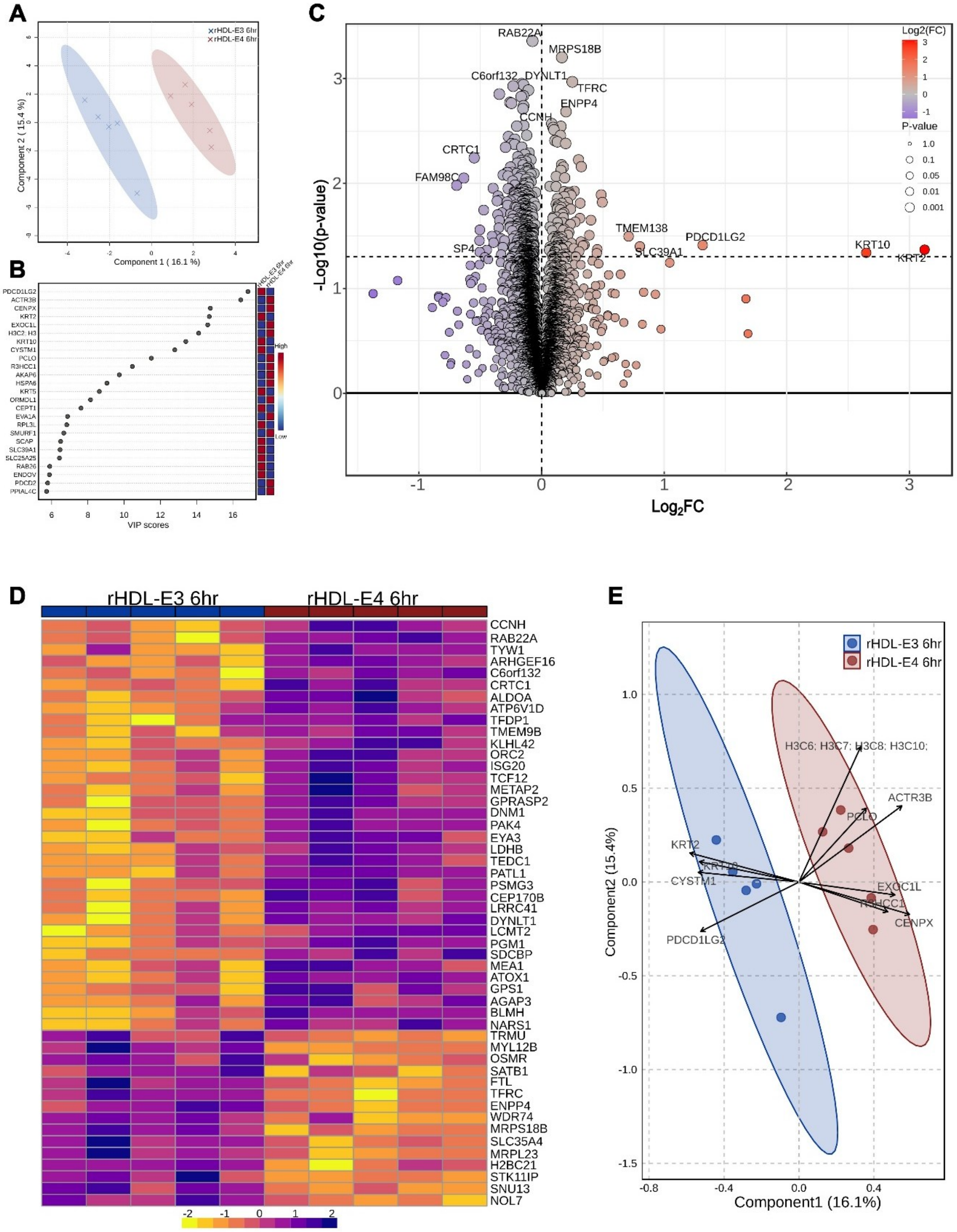


**Supplemental Figure 4. Comparative proteomics of microglia treated with either rHDL-E3 or rHDL-E4 for 6 hours.** **A.** PLS-DA plot of rHDL-E3 vs rHDL-E4 treatment outcomes at the 6 h time point. **B.** VIP score table displaying the top 25 proteins driving differences in the PLS-DA plot. **C.** Volcano plot displaying rHDL-E3 vs. rHDL-E4 with x-axis representing $\log_2$FC and y-axis representing -$\log_{10}$(p-value). Purple indicates downregulated proteins in response to rHDL-E3 and orange indicates upregulated proteins in response to rHDL-E3 treatment, at 6-h. **D.** Heatmap of the top 50 DAP proteins with color gradients associated with the normalized concentrations. **E.** PLS-DA biplot with arrows representing gene loadings; direction and length indicate the contribution of specific proteins to the sample separation. Proteins pointing to a specific cluster are positively associated with that group.

**Supplementary Table 4A.** Upregulated biological pathways in LF-E3 vs rHDL-E3 at 24 hr

| Biological Pathway | Protein count | Background protein count | Strength | Signal | FDR | Proteins |
|---|---|---|---|---|---|---|
| Immune response | 80 | 377 | 0.6 | 1.67 | 3.90e-20 | CTSC,POLR3B,OAS3,EIF2AK2,ZC3HAV1,BST2,SHFL,RFX1,TRIM21,OASL,VAMP7,CD81,HERC5,PML,PTX3,DTX3L,ERAP1,TMEM43,TRIM56,CEBPB,ISG20,APOBEC3F,ADAM17,STAT2,IL17RA,TRIM25,APOBEC3B,SMAD3,MX2,OAS2,SP100,TRIM14,LYAR,TAP1,TRIM38,PARP9,STAT1,DHX9,IFI16,ADAR,NFKB2,GBP1,GBP3,IFI44,ZNFX1,IFIT5,IFIT1,IFIT3,CD40,POLR3A,TAP2,HLA-C,HLA-E,PIK3CD,DDX58,TRIM22,DDX60,LGALS9,RAB27A,HLA-A,IRF7,MX1,OAS1,PRKD2,CD70,IFI35,CHID1,HLA-B,ERAP2,MAVS,TAPBP,PARP14,TREX1,IFIT2,SAMHD1,ISG15,MYO1C,IFIH1,B2M,MYD88 |
| Innate immune response | 65 | 250 | 0.69 | 1.91 | 6.30e-20 | POLR3B,OAS3,EIF2AK2,ZC3HAV1,BST2,SHFL,TRIM21,OASL,VAMP7,HERC5,PML,PTX3,DTX3L,TMEM43,TRIM56,ISG20,APOBEC3F,STAT2,IL17RA,TRIM25,APOBEC3B,MX2,OAS2,SP100,TRIM14,LYAR,TRIM38,PARP9,STAT1,DHX9,IFI16,ADAR,GBP1,GBP3,ZNFX1,IFIT5,IFIT1,IFIT3,CD40,POLR3A,HLA-C,HLA-E,PIK3CD,DDX58,TRIM22,DDX60,LGALS9,RAB27A,HLA-A,IRF7,MX1,OAS1,IFI35,CHID1,HLA-B,MAVS,PARP14,TREX1,IFIT2,SAMHD1,ISG15,MYO1C,IFIH1,B2M,MYD88 |
| Defense response to virus | 46 | 119 | 0.86 | 2.36 | 4.82e-19 | POLR3B,OAS3,EIF2AK2,ZC3HAV1,BST2,SHFL,TRIM21,OASL,HERC5,PML,DTX3L,TRIM56,ISG20,APOBEC3F,STAT2,TRIM25,APOBEC3B,MX2,OAS2,PLSCR1,PARP9,STAT1,IFI16,ADAR,GBP1,GBP3,ZNFX1,IFIT5,IFIT1,IFIT3,CD40,POLR3A,DDX58,TRIM22,DDX60,IRF9,IRF7,MX1,OAS1,MAVS,TREX1,IFIT2,SAMHD1,ISG15,IFIH1,MYD88 |
| Defense response to other organism | 69 | 316 | 0.61 | 1.65 | 4.58e-18 | POLR3B,OAS3,EIF2AK2,ZC3HAV1,BST2,SHFL,TRIM21,OASL,VAMP7,HERC5,PML,PTX3,DTX3L,TMEM43,TRIM56,CEBPB,ISG20,APOBEC3F,ADAM17,STAT2,IL17RA,TRIM25,APOBEC3B,MX2,OAS2,SP100,TRIM14,PLSCR1,LYAR,TRIM38,PARP9,STAT1,DHX9,IFI16,ADAR,GBP1,GBP3,ZNFX1,IFIT5,IFIT1,IFIT3,CD40,POLR3A,HLA-C,HLA-E,PIK3CD,DDX58,TRIM22,DDX60,LGALS9,RAB27A,HLA-A,IRF9,IRF7,MX1,OAS1,IFI35,CHID1,HLA-B,MAVS,PARP14,TREX1,IFIT2,SAMHD1,ISG15,MYO1C,IFIH1,B2M,MYD88 |
| Response to virus | 50 | 173 | 0.73 | 1.89 | 9.49e-17 | POLR3B,OAS3,EIF2AK2,ZC3HAV1,BST2,SHFL,TRIM21,OASL,HERC5,PML,DTX3L,TRIM56,ISG20,APOBEC3F,STAT2,IL17RA,TRIM25,APOBEC3B,SMAD3,MX2,OAS2,PLSCR1,PARP9,STAT1,IFI16,ADAR,GBP1,GBP3,IFI44,ZNFX1,IFIT5,IFIT1,IFIT3,CD40,POLR3A,DDX58,TRIM22,DDX60,LGALS9,IRF9,IRF7,MX1,OAS1,MAVS,TREX1,IFIT2,SAMHD1,ISG15,IFIH1,MYD88 |
| Regulation of viral process | 34 | 100 | 0.8 | 1.79 | 1.00e-12 | OAS3,EIF2AK2,ZC3HAV1,NECTIN2,BST2,SHFL,TRIM21,OASL,HACD3,PTX3,ISG20,APOBEC3F,TRIM25,APOBEC3B,OAS2,SP100,TRIM14,PLSCR1,TRIM38,STAT1,DHX9,IFI16,ADAR,ZNFX1,IFIT5,IFIT1,TRIM22,LGALS9,MX1,OAS1,MAVS,PARP10,ISG15,IFIH1 |
| Negative regulation of viral process | 27 | 57 | 0.95 | 2.04 | 1.39e-12 | OAS3,EIF2AK2,ZC3HAV1,BST2,SHFL,TRIM21,OASL,PTX3,ISG20,APOBEC3F,TRIM25,APOBEC3B,OAS2,SP100,TRIM14,PLSCR1,STAT1,IFI16,ZNFX1,IFIT5,IFIT1,MX1,OAS1,MAVS,PARP10,ISG15,IFIH1 |
| Regulation of viral life cycle | 31 | 83 | 0.84 | 1.84 | 2.03e-12 | OAS3,EIF2AK2,ZC3HAV1,NECTIN2,BST2,SHFL,TRIM21,OASL,HACD3,PTX3,ISG20,APOBEC3F,TRIM25,APOBEC3B,OAS2,TRIM14,PLSCR1,TRIM38,IFI16,ADAR,ZNFX1,IFIT5,IFIT1,TRIM22,LGALS9,MX1,OAS1,MAVS,PARP10,ISG15,IFIH1 |
| Negative regulation of viral genome replication | 21 | 37 | 1.03 | 1.93 | 1.06e-10 | OAS3,EIF2AK2,ZC3HAV1,BST2,SHFL,OASL,ISG20,APOBEC3F,APOBEC3B,OAS2,PLSCR1,IFI16,ZNFX1,IFIT5,IFIT1,MX1,OAS1,MAVS,PARP10,ISG15,IFIH1 |
| Regulation of viral genome replication | 24 | 59 | 0.88 | 1.65 | 5.44e-10 | OAS3,EIF2AK2,ZC3HAV1,BST2,SHFL,OASL,HACD3,ISG20,APOBEC3F,APOBEC3B,OAS2,PLSCR1,TRIM38,IFI16,ADAR,ZNFX1,IFIT5,IFIT1,MX1,OAS1,MAVS,PARP10,ISG15,IFIH1 |

**Supplementary Table 4B.** Downregulated biological pathways in LF-E3 vs rHDL-E3 at 24 hr

| Biological Pathway | Protein count | Background protein count | Strength | Signal | FDR | Protein |
|---|---|---|---|---|---|---|
| Positive regulation of protein catabolic process | 21 | 116 | 0.58 | 0.47 | 0.0075 | USP5,APOE,SNX1,USP13,SH3RF1,APP,DDB1,PSMC5,DAB2,RNF14,EZR,VCP,BAG2,HSPA1A,BAG6,KEAP1,HECTD1,PSMC2,PSMC3,PSMC6,RDX |
| Regulation of protein catabolic process | 28 | 216 | 0.44 | 0.35 | 0.0232 | USP5,APOE,SNX1,USP13,SH3RF1,APP,DDB1,PSMC5,DAB2,FMN2,NQO1,RNF14,HGS,PSMF1,EZR,ANXA2,VCP,BAG2,HSPA1A,BAG6,KEAP1,HECTD1,PSMC2,PSMC3,SNX12,FAM83D,PSMC6,RDX |
| Positive regulation of proteasomal protein catabolic process | 15 | 72 | 0.64 | 0.39 | 0.0232 | USP5,USP13,SH3RF1,PSMC5,DAB2,RNF14,VCP,BAG2,HSPA1A,BAG6,KEAP1,HECTD1,PSMC2,PSMC3,PSMC6 |
| Regulation of catabolic process | 53 | 596 | 0.27 | 0.3 | 0.0345 | PIK3R2,APOC3,USP5,FXR2,ZC3H14,APOE,FUS,TTC5,SNX1,PIK3C3,USP13,CASC3,ATP6V1A,ATP6V1B2,SH3RF1,APP,WDR41,DDB1,PSMC5,DAB2,FMN2,NQO1,RNF14,HGS,PSMF1,EZR,ATG101,ANXA2,CNOT3,VCP,EXOC8,BAG2,HSPA1A,BAG6,CTTN,SNX5,KEAP1,CARHSP1,GAPDH,KHSRP,HECTD1,MTMR3,PSMC2,TNRC6B,IGF2BP2,NRBP2,ATG13,PSMC3,SNX12,FAM83D,CSDE1,PSMC6,RDX |
| Regulation of vesicle-mediated transport | 30 | 259 | 0.39 | 0.32 | 0.0345 | APOC3,SNX17,TSG101,APOE,PACSIN2,PTPN23,ATG3,APP,WDR41,AXL,FGB,DAB2,MKLN1,STXBP6,RUFY1,HGS,FGG,EZR,ANXA2,S100A10,SDC4,HNRNPK,ANXA1,STAM,KIF3A,PDCD6IP,ATXN2,CNN2,SNX12,RDX |

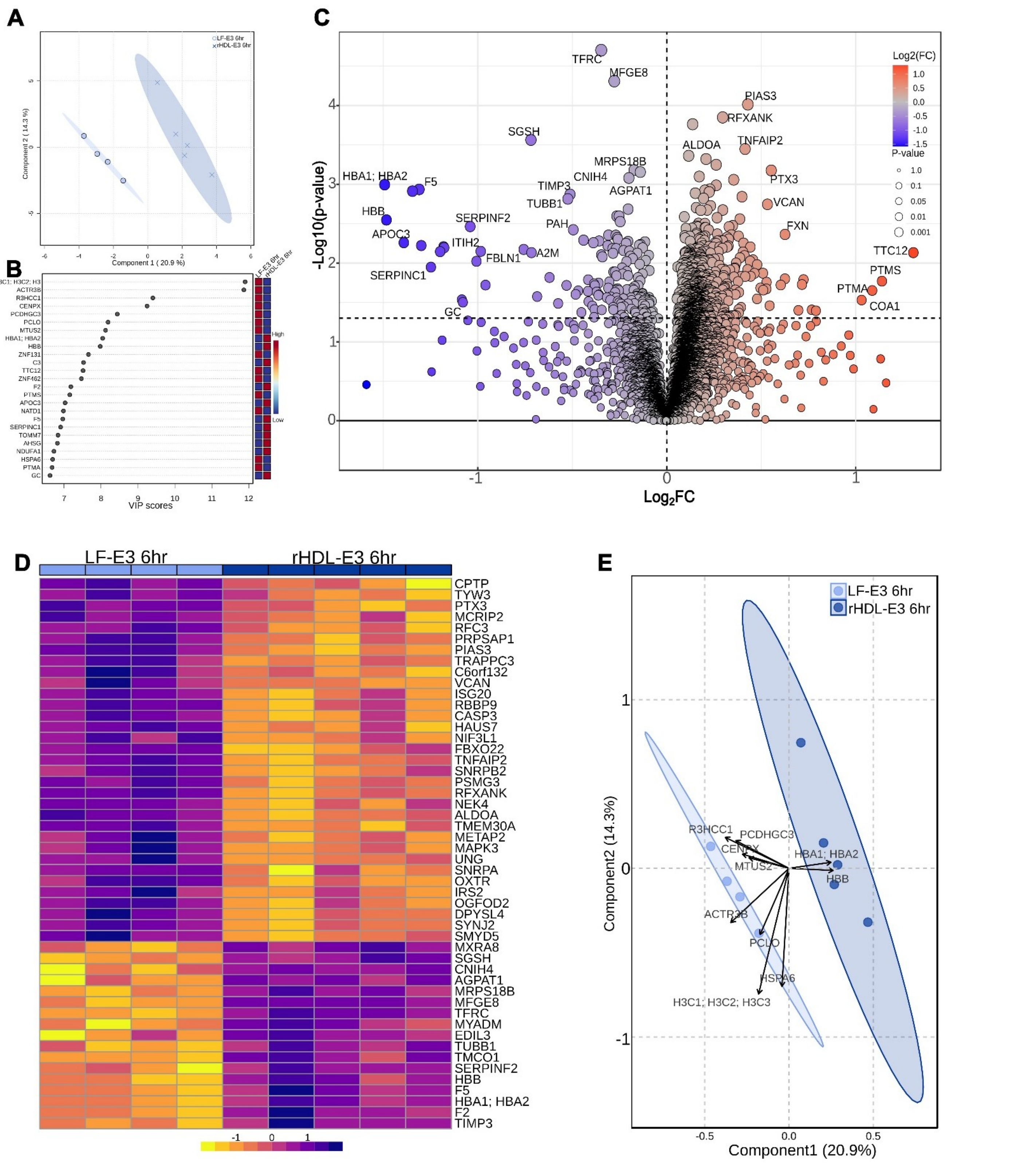

**Supplemental Figure 5. Comparative proteomics of microglia treated with either LF-E3 or rHDL-E3 for 6 hours**. **A.** PLS-DA plot of LF-E3 vs rHDL-E3 treatment outcomes at the 6 h time point. **B.** VIP score table displaying the top 25 proteins driving differences in the PLS-DA plot. **C.** Volcano plot displaying LF-E3 with x-axis representing $log_2FC$ and y-axis representing $-log_{10}$(p-value). Purple indicates downregulated proteins in response to LF-E3 and orange indicates upregulated proteins in response to LF-E3 treatment, at 6 h. **D.** Heatmap of the top 50 DAP proteins with color gradients associated with the normalized concentrations. **E**. PLS-DA biplot with arrows representing gene loadings; direction and length indicate the contribution of specific proteins to the sample separation. Proteins pointing to a specific cluster are positively associated with that group.

**Supplementary Table 5A.** Upregulated biological pathways in LF-E4 vs rHDL-E4 at 24 hr

| Biological Pathway | Protein count | Background protein count | Strength | Signal | FDR | Proteins |
|---|---|---|---|---|---|---|
| Defense response | 60 | 426 | 0.89 | 2.77 | 5.23e-33 | NFKB1,OAS3,EIF2AK2,ZC3HAV1,NMI,APOL2,BST2,TRIM21,OASL,LGALS3BP,LOXL3,HERC5,PML,PTX3,DTX3L,ISG20,STAT2,TRIM25,MX2,LIPA,OAS2,SP100,TRIM14,PLSCR1,TAP1,RBCK1,TRIM38,PARP9,STAT1,ADAR,GBP1,GBP3,ZNFX1,IFIT5,IFIT1,IFIT3,HLA-C,HLA-E,PIK3CD,DDX58,TRIM22,TRIM5,PARP4,TFRC,DDX60,LGALS9,IRF9,IRF7,MX1,OAS1,IFI35,HLA-B,PARP14,LGALS8,IFIT2,SAMHD1,ISG15,IFIH1,B2M,MYD88 |
| Response to virus | 43 | 173 | 1.14 | 3.89 | 2.05e-31 | NFKB1,OAS3,EIF2AK2,ZC3HAV1,NMI,BST2,TRIM21,OASL,HERC5,PML,DTX3L,ISG20,STAT2,TRIM25,MX2,OAS2,PLSCR1,PARP9,STAT1,ADAR,GBP1,FMR1,GBP3,IFI44,ZNFX1,IFIT5,IFIT1,IFIT3,DDX58,TRIM22,TRIM5,DDX60,LGALS9,IRF9,IRF7,MX1,OAS1,LGALS8,IFIT2,SAMHD1,ISG15,IFIH1,MYD88 |
| Defense response to other organism | 52 | 316 | 0.96 | 3.02 | 2.39e-31 | OAS3,EIF2AK2,ZC3HAV1,NMI,BST2,TRIM21,OASL,HERC5,PML,PTX3,DTX3L,ISG20,STAT2,TRIM25,MX2,OAS2,SP100,TRIM14,PLSCR1,RBCK1,TRIM38,PARP9,STAT1,ADAR,GBP1,GBP3,ZNFX1,IFIT5,IFIT1,IFIT3,HLA-C,HLA-E,PIK3CD,DDX58,TRIM22,TRIM5,DDX60,LGALS9,IRF9,IRF7,MX1,OAS1,IFI35,HLA-B,PARP14,LGALS8,IFIT2,SAMHD1,ISG15,IFIH1,B2M,MYD88 |
| Innate immune response | 48 | 250 | 1.03 | 3.33 | 2.90e-31 | OAS3,EIF2AK2,ZC3HAV1,NMI,BST2,TRIM21,OASL,HERC5,PML,PTX3,DTX3L,ISG20,STAT2,TRIM25,MX2,OAS2,SP100,TRIM14,TRIM38,PARP9,STAT1,ADAR,GBP1,GBP3,ZNFX1,IFIT5,IFIT1,IFIT3,HLA-C,HLA-E,PIK3CD,DDX58,TRIM22,TRIM5,DDX60,LGALS9,IRF7,MX1,OAS1,IFI35,HLA-B,PARP14,IFIT2,SAMHD1,ISG15,IFIH1,B2M,MYD88 |
| Immune response | 55 | 377 | 0.91 | 2.78 | 3.36e-31 | OAS3,EIF2AK2,ZC3HAV1,NMI,BST2,TRIM21,OASL,HERC5,PML,PTX3,DTX3L,ERAP1,ISG20,STAT2,TRIM25,MX2,OAS2,SP100,TRIM14,TAP1,TRIM38,PSMB10,PARP9,STAT1,ADAR,NFKB2,GBP1,GBP3,IFI44,ZNFX1,IFIT5,IFIT1,IFIT3,HLA-C,HLA-E,PIK3CD,DDX58,TRIM22,TRIM5,DDX60,LGALS9,IRF7,MX1,OAS1,IFI35,HLA-B,ERAP2,TAPBP,PARP14,IFIT2,SAMHD1,ISG15,IFIH1,B2M,MYD88 |
| Defense response to virus | 37 | 119 | 1.24 | 4.25 | 1.28e-29 | OAS3,EIF2AK2,ZC3HAV1,BST2,TRIM21,OASL,HERC5,PML,DTX3L,ISG20,STAT2,TRIM25,MX2,OAS2,PLSCR1,PARP9,STAT1,ADAR,GBP1,GBP3,ZNFX1,IFIT5,IFIT1,IFIT3,DDX58,TRIM22,TRIM5,DDX60,IRF9,IRF7,MX1,OAS1,IFIT2,SAMHD1,ISG15,IFIH1,MYD88 |
| Regulation of viral process | 29 | 100 | 1.2 | 3.48 | 6.45e-22 | OAS3,EIF2AK2,ZC3HAV1,NECTIN2,BST2,TRIM21,OASL,PTX3,ISG20,TRIM25,OAS2,SP100,TRIM14,PLSCR1,TRIM38,STAT1,ADAR,FMR1,ZNFX1,IFIT5,IFIT1,TRIM22,TRIM5,LGALS9,MX1,OAS1,PARP10,ISG15,IFIH1 |
| Regulation of viral life cycle | 27 | 83 | 1.25 | 3.59 | 2.32e-21 | OAS3,EIF2AK2,ZC3HAV1,NECTIN2,BST2,TRIM21,OASL,PTX3,ISG20,TRIM25,OAS2,TRIM14,PLSCR1,TRIM38,ADAR,FMR1,ZNFX1,IFIT5,IFIT1,TRIM22,TRIM5,LGALS9,MX1,OAS1,PARP10,ISG15,IFIH1 |
| Negative regulation of viral process | 23 | 57 | 1.35 | 3.66 | 1.29e-19 | OAS3,EIF2AK2,ZC3HAV1,BST2,TRIM21,OASL,PTX3,ISG20,TRIM25,OAS2,SP100,TRIM14,PLSCR1,STAT1,ZNFX1,IFIT5,IFIT1,TRIM5,MX1,OAS1,PARP10,ISG15,IFIH1 |
| Negative regulation of viral genome replication | 16 | 37 | 1.38 | 2.83 | 1.35e-13 | OAS3,EIF2AK2,ZC3HAV1,BST2,OASL,ISG20,OAS2,PLSCR1,ZNFX1,IFIT5,IFIT1,MX1,OAS1,PARP10,ISG15,IFIH1 |

**Supplementary Table 5B.** Upregulated KEGG pathways in LF-E4 vs rHDL-E4 at 24 hr

| KEGG Pathway | Protein count | Background protein count | Strengt h | Signal | FDR | Protein |
|---|---|---|---|---|---|---|
| Herpes simplex virus 1 infection | 23 | 102 | 1.1 | 2.5 | 4.17e-15 | NFKB1,OAS3,EIF2AK2,BST2,PML,ITGA5,STAT2,OAS2,SP100,TAP1,STAT1,HLA-C,HLA-E,PIK3CD,DDX58,IRF9,IRF7,OAS1,HLA-B,TAPBP,IFIH1,B2M,MYD88 |
| Influenza A | 18 | 79 | 1.1 | 2.12 | 8.00e-12 | NFKB1,OAS3,EIF2AK2,PML,STAT2,TRIM25,MX2,OAS2,STAT1,ADAR,PIK3CD,DDX58,IRF9,IRF7,MX1,OAS1,IFIH1,MYD88 |
| Epstein-Barr virus infection | 20 | 115 | 0.98 | 1.89 | 1.68e-11 | NFKB1,OAS3,EIF2AK2,STAT2,OAS2,TAP1,STAT1,NFKB2,HLA-C,HLA-E,PIK3CD,DDX58,IRF9,IRF7,OAS1,HLA-B,TAPBP,ISG15,B2M,MYD88 |
| Measles | 16 | 78 | 1.05 | 1.83 | 4.54e-10 | NFKB1,OAS3,EIF2AK2,STAT2,MX2,OAS2,STAT1,ADAR,PIK3CD,DDX58,IRF9,IRF7,MX1,OAS1,IFIH1,MYD88 |
| Hepatitis C | 14 | 87 | 0.95 | 1.37 | 1.41e-07 | NFKB1,OAS3,EIF2AK2,STAT2,MX2,OAS2,STAT1,IFIT1,PIK3CD,DDX58,IRF9,IRF7,MX1,OAS1 |
| Human papillomavirus infection | 17 | 165 | 0.76 | 1.08 | 9.75e-07 | NFKB1,EIF2AK2,OASL,ITGAV,ITGA5,COL6A2,STAT2,MX2,STAT1,COL6A1,HLA-C,HLA-E,PIK3CD,IRF9,MX1,HLA-B,ISG15 |
| NOD-like receptor signaling pathway | 12 | 86 | 0.89 | 1.08 | 6.21e-06 | NFKB1,OAS3,STAT2,OAS2,RBCK1,STAT1,GBP1,GBP3,IRF9,IRF7,OAS1,MYD88 |
| Viral carcinogenesis | 12 | 118 | 0.75 | 0.81 | 0.00011 | NFKB1,EIF2AK2,SP100,NFKB2,GTF2B,HLA-C,HLA-E,PIK3CD,IL6ST,IRF9,IRF7,HLA-B |
| Kaposi sarcoma-associated herpesvirus infection | 11 | 96 | 0.8 | 0.84 | 0.00011 | NFKB1,EIF2AK2,STAT2,STAT1,HLA-C,HLA-E,PIK3CD,IL6ST,IRF9,IRF7,HLA-B |
| Human immunodeficiency virus 1 infection | 12 | 129 | 0.71 | 0.75 | 0.00022 | NFKB1,BST2,TAP1,HLA-C,HLA-E,PIK3CD,TRIM5,HLA-B,TAPBP,SAMHD1,B2M,MYD88 |

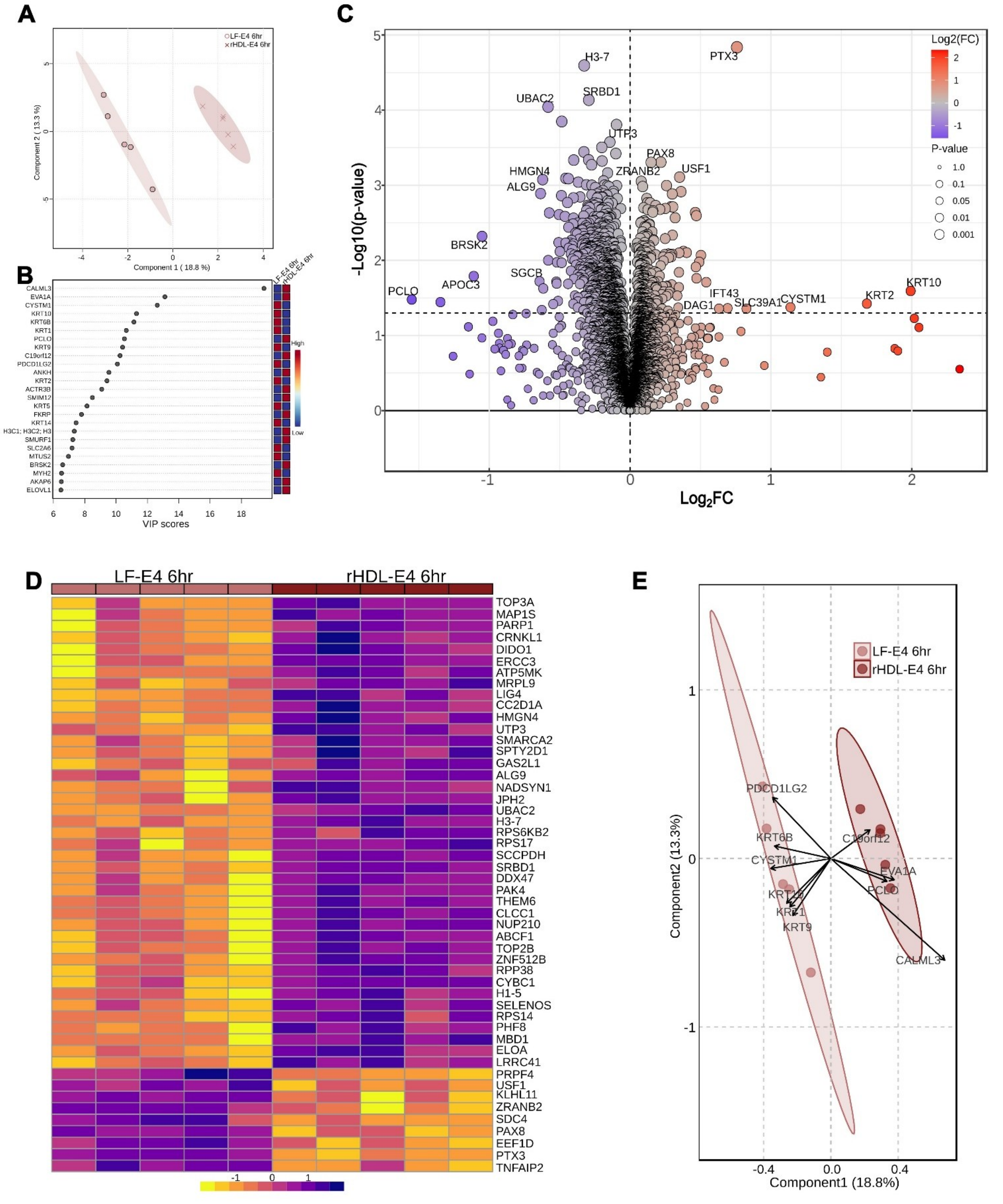


**Supplemental Figure 6**. **Comparative proteomics of microglia treated with either LF-E4 or rHDL-E4 for 6 hours. A.** PLS-DA plot of LF-E4 vs rHDL-E4 treatment outcomes at the 6 h time point. **B.** VIP score table displaying the top 25 proteins driving differences in the PLS-DA plot. **C.** Volcano plot displaying LF-E4 vs rHDL-E4 with x-axis representing $log_2FC$ and y-axis representing $-log_{10}$(p-value). Purple indicates downregulated proteins in response to LF-E4 and orange indicates upregulated proteins in response to LF-E4 treatment, at 6 h. **D.** Heatmap of the top 50 DAP proteins with color gradients associated with the normalized concentrations. **E**. PLS-DA biplot with arrows representing gene loadings; direction and length indicate the contribution of specific proteins to the sample separation. Proteins pointing to a specific cluster are positively associated with that group.

**Supplemental Table 6A.** Upregulated KEGG pathways in LF-E3 vs Ctrl at 24 h

| KEGG Pathway | Protein count | Background protein count | Strength | Signal | FDR | Protein |
|---|---|---|---|---|---|---|
| Influenza A | 17 | 79 | 0.61 | 0.58 | 0.0017 | PML,BAX,STAT2,TRIM25,MX2,TRADD,OAS2,FDPS,STAT1,DDX58,IRF9,IRF7,MX1,RELA,OAS1,IFIH1,MYD88 |
| Hepatitis C | 17 | 87 | 0.56 | 0.55 | 0.0021 | STAT3,BAX,STAT2,PPP2R2A,MX2,TRADD,OAS2,STAT1,IFIT1,YWHAB,DDX58,IRF9,IRF7,MX1,RELA,OAS1,LDLR |
| Epstein-Barr virus infection | 20 | 115 | 0.51 | 0.54 | 0.0021 | STAT3,BAX,STAT2,CALR,TRADD,OAS2,STAT1,NFKB2,HLA-C,DDX58,IRF9,IRF7,RELA,PSMD14,OAS1,CD44,HLA-B,LYN,ISG15,MYD88 |
| Measles | 15 | 78 | 0.56 | 0.5 | 0.0048 | STAT3,BAX,STAT2,MX2,TRADD,OAS2,STAT1,DDX58,IRF9,IRF7,MX1,RELA,OAS1,IFIH1,MYD88 |
| Herpes simplex virus 1 infection | 17 | 102 | 0.5 | 0.46 | 0.0061 | PML,BAX,STAT2,CALR,TRADD,OAS2,SP100,STAT1,HLA-C,DDX58,IRF9,IRF7,RELA,OAS1,HLA-B,IFIH1,MYD88 |
| RIG-I-like receptor signaling pathway | 9 | 35 | 0.68 | 0.42 | 0.0159 | OTUD5,TRIM25,TRADD,DDX58,IRF7,RELA,CYLD,ISG15,IFIH1 |
| Terpenoid backbone biosynthesis | 6 | 15 | 0.88 | 0.42 | 0.0216 | MVK,HMGCR,HMGCS1,FDPS,ACAT2,IDI1 |

**Supplemental Table 6B.** Upregulated KEGG pathways in LF-E4 vs Ctrl at 24 h

| KEGG Pathway | Protein count | Background protein count | Strength | Signal | FDR | Protein |
|---|---|---|---|---|---|---|
| Influenza A | 16 | 79 | 0.88 | 1.22 | 7.35e-07 | OAS3,PML,STAT2,TRIM25,MX2,OAS2,FDPS,STAT1,ADAR,DDX58,IRF9,IRF7,MX1,OAS1,IFIH1,MYD88 |
| Measles | 14 | 78 | 0.83 | 0.99 | 1.47e-05 | OAS3,STAT3,STAT2,MX2,OAS2,STAT1,ADAR,DDX58,IRF9,IRF7,MX1,OAS1,IFIH1,MYD88 |
| Herpes simplex virus 1 infection | 15 | 102 | 0.74 | 0.9 | 2.70e-05 | OAS3,PML,STAT2,CALR,OAS2,SP100,STAT1,HLA-C,DDX58,IRF9,IRF7,OAS1,HLA-B,IFIH1,MYD88 |
| Epstein-Barr virus infection | 16 | 115 | 0.72 | 0.89 | 2.70e-05 | OAS3,STAT3,STAT2,CALR,OAS2,STAT1,NFKB2,HLA-C,DDX58,IRF9,IRF7,NCOR2,OAS1,HLA-B,ISG15,MYD88 |
| Terpenoid backbone biosynthesis | 7 | 15 | 1.24 | 1.06 | 5.95e-05 | MVK,HMGCR,MVD,HMGCS1,FDPS,ACAT2,IDI1 |
| Hepatitis C | 13 | 87 | 0.75 | 0.83 | 9.23e-05 | OAS3,STAT3,STAT2,MX2,OAS2,STAT1,IFIT1,DDX58,IRF9,IRF7,MX1,OAS1,LDLR |
| Steroid biosynthesis | 6 | 13 | 1.24 | 0.91 | 0.00028 | CYP51A1,MSMO1,SQLE,DHCR7,LSS,FDFT1 |
| NOD-like receptor signaling pathway | 11 | 86 | 0.68 | 0.6 | 0.0017 | GABARAPL2,OAS3,STAT2,OAS2,STAT1,GBP1,IRF9,IRF7,OAS1,ERBIN,MYD88 |
| Metabolic pathways | 36 | 734 | 0.26 | 0.35 | 0.0123 | CYP51A1,PCK2,LAP3,MVK,MSMO1,SQLE,FADS2,PLOD2,HMGCR,MVD,FASN,ASL,GLS,HMGCS1,ACSL4,GPX4,DHCR7,FDPS,ACAT2,SCD,NEU1,PSAT1,IDI1,ASNS,LSS,CBS,FAH,UAP1L1,PLD3,IDH1,PCYT2,ALDH6A1,HEXA,ACACA,FDFT1,PHGDH |

| | | | | | | |
|---|---|---|---|---|---|---|
| Cell adhesion molecules | 6 | 31 | 0.86 | 0.47 | 0.0123 | ITGAV,ALCAM,PTPRF,L1CAM,HLA-C,HLA-B |
| RIG-I-like receptor signaling pathway | 6 | 35 | 0.81 | 0.43 | 0.0184 | TRIM25,DDX58,IRF7,CYLD,ISG15,IFIH1 |
| Human papillomavirus infection | 13 | 165 | 0.47 | 0.37 | 0.0202 | OASL,ITGAV,COL6A2,ITGA3,STAT2,MX2,STAT1,COL6A1,HLA-C,IRF9,MX1,HLA-B,ISG15 |
| Fatty acid metabolism | 6 | 39 | 0.76 | 0.39 | 0.0258 | FADS2,FASN,ACSL4,ACAT2,SCD,ACACA |
| PPAR signaling pathway | 5 | 26 | 0.86 | 0.4 | 0.0264 | PCK2,FADS2,HMGCS1,ACSL4,SCD |
